# RNA quality control by CCR4 safeguards chromatin integrity and centromere function in *Arabidopsis*

**DOI:** 10.1101/2024.08.23.608929

**Authors:** Atsushi Shimada, Hidetoshi Saze

## Abstract

The centromere is a pivotal chromatin domain that ensures accurate chromosome segregation during cell division. However, the epigenome regulation of the centromere and its impact on centromere function remain largely elusive. Here in the model plant *Arabidopsis*, we show that CCR4, the catalytic subunit of the RNA deadenylation complex CCR4-NOT, is essential for maintenance of the centromere epigenome and chromosome integrity. We demonstrate that CCR4 is involved in shortening of the poly(A) tails of transcripts originated from centromeric transposons and repeats, thereby promoting the production of small interfering RNAs (siRNAs). The CCR4-dependent siRNAs guide non-CG DNA methylation at centromere repeats, and CCR4 cooperates with canonical DNA methylation pathways to enhance centromeric H3K9 methylation and ensure mitotic chromosome stability. Our study illustrates the crucial role of RNA quality control in RNA interference and reveals the elaborate mechanism that safeguards plant centromeres through epigenomic regulation.

## Introduction

In eukaryotes, the centromere plays a crucial role in ensuring faithful chromosome segregation through mitosis and meiosis. Centromeres in eukaryotes generally contain repetitive sequences, such as short satellite repeats and transposable elements (TEs). These satellite repeats are bound by centromere-specific histone variants that assemble into a complex protein structure called the kinetochore, which serves as a scaffold for spindle microtubule attachment during cell division^1^. Impaired chromatin organization at the centromere leads to chromosome instability and karyotype changes, profoundly impacting cellular traits and genome evolution in both animals and plants ^2, 3^. Although comprehensive analysis of centromeric chromatin has been challenging due to its complex repetitive structures, recent advancements in long-read sequencing technologies have enabled continuous sequence assembly of centromeric regions of chromosomes^4–7^, which are expected to significantly enhance our understanding of chromatin regulation at the centromere. Recent reports on the complete Telomere-to-Telomere (T2T) genome assembly of *Arabidopsis thaliana* revealed the genetic and epigenetic landscape of the *Arabidopsis* centromeres^6, 7^. Similar to other eukaryotic centromeres, the *Arabidopsis* centromeres consist of 178 base pair (bp) satellite repeats and TEs^6, 8^. The core centromere 178 bp repeats are bound by the centromere-specific histone H3 variant (CenH3)^1^. The CenH3-bound centromere is embedded in transcriptionally inactive, densely packed chromatin domains called pericentromeric heterochromatin. This heterochromatin primarily consists of satellite repeats and degenerated TE sequences and is associated with repressive chromatin modifications such as DNA methylation and histone H3K9 methylation^6^. Studies in yeast have shown that pericentromeric heterochromatin is required for sister chromatid cohesion during mitosis^9^.

Plants utilize RNA interference (RNAi) mechanism to guide DNA methylation and induce heterochromatin formation, a process called RNA-directed DNA methylation (RdDM)^10^. In the RdDM pathway, TE and repeat-derived RNAs transcribed by plant-specific RNA polymerase V are converted to double-stranded RNA (dsRNA) by RNA-dependent RNA polymerase 2 (RDR2), which are then cleaved into 24-nucleotide (nt) small interfering RNAs (siRNAs) by DCL3. These siRNAs are loaded onto Argonaute proteins^10^, which recruit *de novo* DNA methyltransferases to target TEs and repeats through siRNA sequence homology^10^. Additionally, a non-canonical RdDM pathway involving RDR6 mediates the production of dsRNA and 21-nt siRNA from RNA polymerase II-derived mRNA^10^. Established heterochromatin is maintained by the chromatin remodeler Decrease in DNA Methylation 1 (DDM1), which allows maintenance DNA methyltransferases to access the heterochromatin^11^. Loss of DDM1 results in a genome-wide reduction in DNA methylation and H3K9 methylation, leading to transcriptional activation of TEs^12^.

Besides RNAi, eukaryotes have evolved an additional RNA quality control factor called CCR4-NOT, which is conserved from yeasts to mammals. CCR4-NOT is a multi-functional protein complex that regulates mRNA biogenesis, from transcription to degradation^13, 14^. The complex is composed of several protein modules with distinct functions, including RNA deadenylases CCR4 and CAF1, which facilitates RNA degradation and turnover by shortening the poly(A) tail of RNA molecules^14^. The *Arabidopsis* CCR4-NOT complex consists of subunits conserved in other organisms^15^. *Arabidopsis* CCR4 (AtCCR4) is localized at cytoplasmic processing bodies (P-bodies), granule structures involved in RNA processing events^16^. Recent studies have shown that AtCCR4 regulates cytoplasmic TE transcripts largely independent of RNAi pathway^17^. However, AtCCR4 is also known to be localized in the nucleus^16^. While CCR4 has been reported to contribute to heterochromatin maintenance in yeast^18, 19^, the function of *Arabidopsis* AtCCR4 in the nucleus, particularly in chromatin regulation, remains poorly understood.

An intriguing mystery in *Arabidopsis* centromere regulation is that mutants defective in DNA methylation pathways do not exhibit severe chromosome instability, despite a significant loss of heterochromatin in pericentromeric regions^20^. This observation suggests the presence of unidentified mechanisms that safeguard centromere function in the absence of primary machinery for heterochromatin maintenance. In this study, we demonstrate that AtCCR4 is a factor that controls heterochromatin and centromere function in synergy with other epigenetic machineries. We show that AtCCR4 promotes RDR6-dependent siRNA synthesis by deadenylating the poly(A) tail of transcripts generated from TEs and centromeric repeats. Further analyses provide compelling evidence that CCR4-dependent siRNAs shape the epigenome of centromere repeats, collaborating with other DNA methylation pathways to maintain centromere chromatin structure and mitotic chromosome stability. Thus, our study provides novel insights into the epigenome regulation of *Arabidopsis* centromeres.

## Results

### Genetic interaction between CCR4-NOT and DNA methylation pathways

To explore the interplay between CCR4 and epigenetic pathways, we first generated *Arabidopsis* mutants of *CCR4A* and *CCR4B*, the *CCR4* homologs identified as components of the *Arabidopsis* CCR4-NOT complex^15^. *CCR4B* was knocked down using an artificial short hairpin in the *ccr4a* T-DNA knock-out line (Figure S1). We then introduced *ccr4a ccr4b* into *ddm1 rdr2* double mutant, which is defective in maintenance and *de novo* DNA methylation pathways. We found that the quadruple mutant *ccr4a ccr4b ddm1 rdr2* (hereafter referred to as *ccdr*), defective in both CCR4 activity and DNA methylation, exhibited more severe growth defects and reduced fertility with abnormal flower development compared to *ccr4a ccr4b* and *ddm1 rdr2* (Figures 1A-1C). These developmental defects were not observed in the segregating siblings of *ccr4a ccr4b ddm1* and *ccr4a ccr4b rdr2 (*Figure S2), suggesting that CCR4 and DNA methylation pathways cooperatively regulate proper *Arabidopsis* development and reproduction.

**Figure 1.**
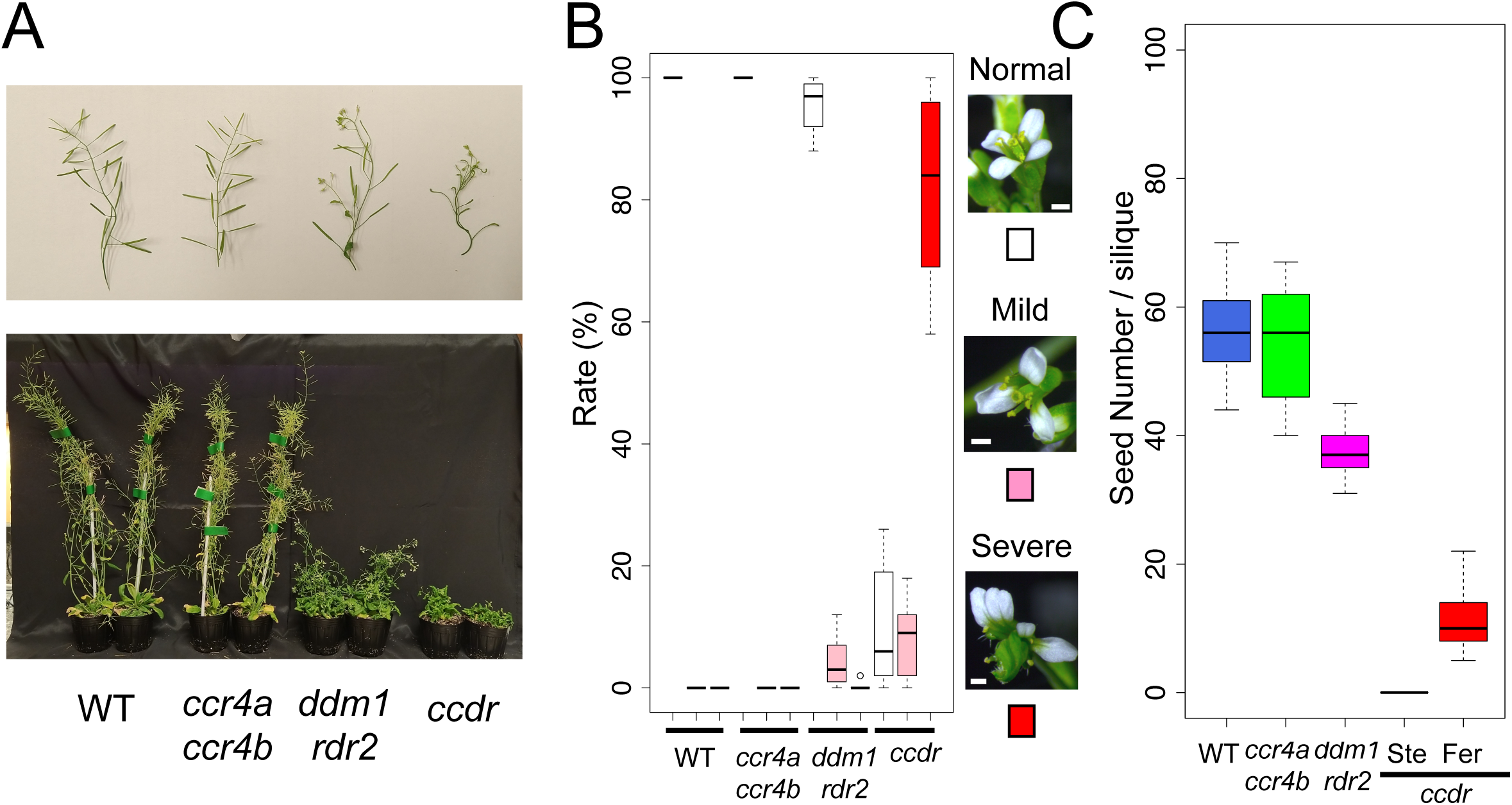
Genetic interaction between CCR4-NOT and DNA methylation pathways. (A) Images of siliques (upper panel) and whole plants (lower panel) of the indicated genotypes. For *ccr4a ccr4b*, *ddm1 rdr2*, and *ccr4a ccr4b ddm1 rdr2* (*ccdr*), the first generation of *ddm1* and/or T4 *ccr4b* knockdown mutants are shown. (B) Boxplots showing flower abnormality in the *ccdr* mutant. Flowers were categorized as Normal (white), Mildly Defective (Mild; pink), or Severely Defective (Severe; red). The percentage of flowers in each category from a single plant is plotted. Fifty flowers were analyzed per plant, with a total of 600 flowers analyzed across 12 plants for each line. The mutant plants were the first generation of *ddm1* and T4 *ccr4b* knockdown lines. (C) Boxplot showing the number of viable seeds in a single silique. Five siliques were analyzed per plant, resulting in a total of 20 siliques analyzed across four plants for each line. Among the 12 *ccdr* plants examined, four produced a few shoots bearing fertile siliques (Fer), while the remaining eight were sterile (Ste). The mutant plants were the first generation of *ddm1* and T4 *ccr4b* knockdown lines.

### CCR4 promotes RDR6-dependent siRNA biogenesis

The results above led us to hypothesize that CCR4 is involved in poly(A) tail processing within epigenetic pathways such as RNAi. To test this, we first assessed the impact of *ccr4* mutations on siRNA biogenesis. Small RNA-seq analysis revealed that, although overall levels of 21-nt small RNA were mostly unaffected in *ccr4a ccr4b* (Figure 2A), several gene loci exhibited reduced levels of 21-nt siRNAs, including the *TAS* loci that produce trans-acting siRNAs (tasiRNAs) (Figure 2B)^21^. We found that all *TAS* genes showed reduced 21-nt siRNA levels in *ccr4a ccr4b* (Figures 2B, 2C and 2E and Table S1). In contrast, the level of microRNAs (miRNAs), another type of 21-nt siRNA, were comparable to those in Wild Type (WT) (Figures 2D and 2E). Since tasiRNAs are known to be produced by the RDR6-dependent RNAi pathway and the poly(A) tail in template RNA inhibits dsRNA synthesis by RDR6 *in vitro*^21–23^, we speculated that CCR4-mediated poly(A) tail shortening promotes RDR6-dependent siRNA synthesis. Indeed, we found that RDR6-dependent 21-nt siRNAs, known as epigenetically activated siRNAs (easiRNAs) ^24^, which accumulate in *ddm1 rdr2*, were absent in *ccdr* (Figure S3A). The representative 21-nt easiRNA-producing *ATHILA* retrotransposons, located in the centromeric region of *Arabidopsis* chromosomes^24^, exhibited a severe decrease in 21-nt siRNAs in *ccdr* (Figures S3B-S3D). The TEs affected in *ccdr* largely overlapped with those affected in *ddm1 rdr2 rdr6* (Figures. S3B, S3E, S3F and S4A). RNA expression levels of genes involved in the RDR6-dependent RNAi pathway were mostly unchanged between *ccdr* and *ddm1 rdr2* (Table S2). Additionally, we did not observe significant changes in TE expressions including easiRNA-associated TEs in *ccdr* compared to *ddm1 rdr2* (Figure S5), suggesting that the loss of easiRNA in *ccdr* is not due to reduced substrate RNAs produced by TEs. Notably, *ddm1 rdr2 rdr6* plants exhibited developmental defects similar to those observed in *ccdr* (Figure S6)^24, 25^. These data indicate that CCR4 activity is required for RDR6-dependent 21-nt siRNA production.

**Figure 2.**
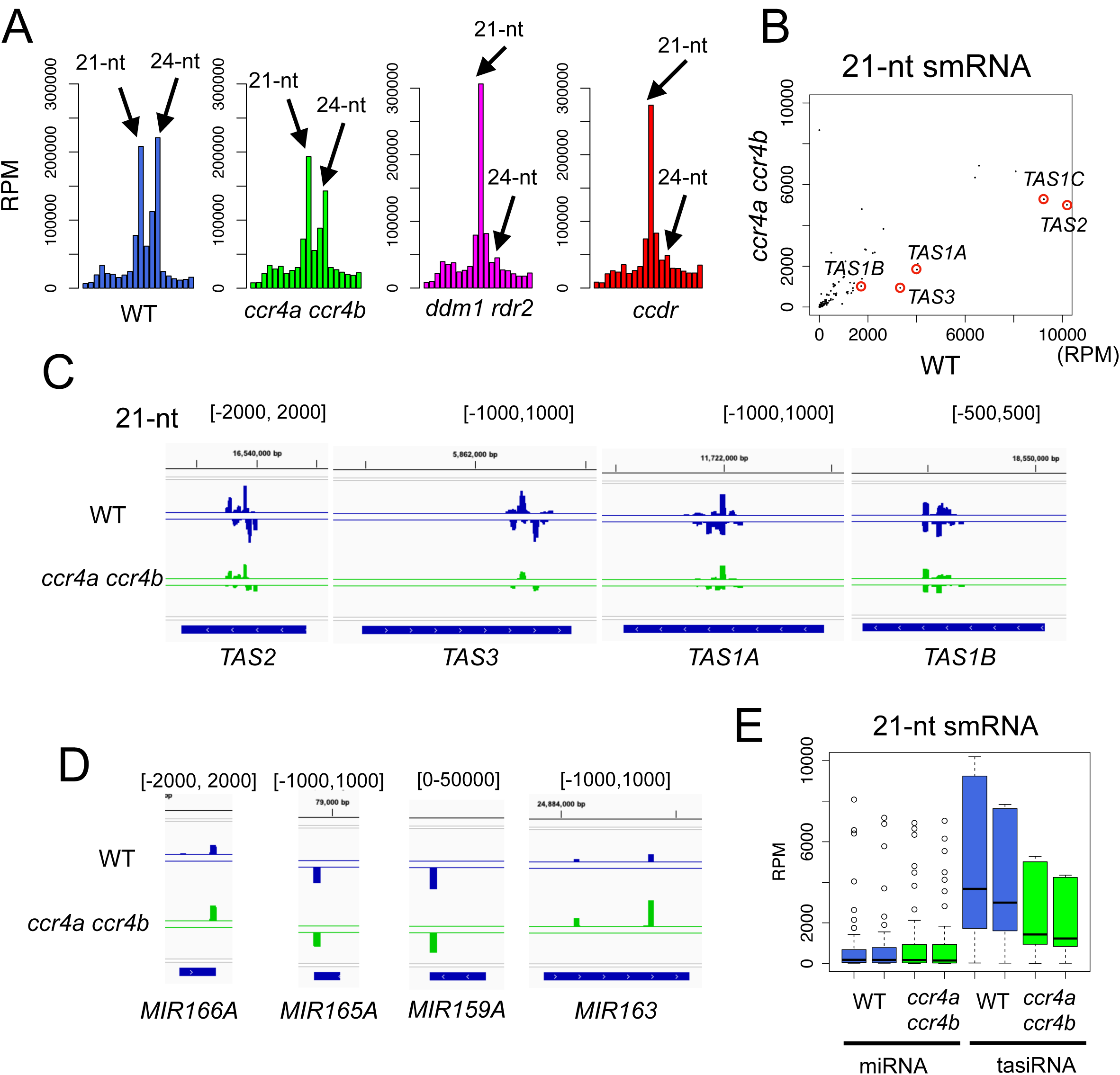
CCR4 is required for 21-nt small RNA biogenesis. (A) Size distribution of small RNAs in the indicated mutant lines. The bar plots show normalized read counts (Reads per Million, RPM) from 11-nt to 30-nt small RNAs. (B) Scatterplot comparing normalized 21-nt small RNA levels (RPM) among genes. 21-nt small RNAs (smRNAs) derived from TAS loci are marked by red circles. (C-D) Normalized 21-nt tasiRNA levels (C) and miRNA levels (D) (RPM) in WT and *ccr4a ccr4b*. (E) Boxplot showing 21-nt tasiRNA and miRNA levels mapped to miRNA genes and TAS loci in WT and *ccr4a ccr4b*.

On the other hand, dsRNA produced by RDR6 is also cleaved into 24-nt siRNAs, which trigger DNA methylation through a non-canonical RdDM pathway^26^. We investigated whether CCR4 also regulates RDR6-dependent 24-nt siRNA synthesis. Small RNA sequencing data showed that, similar to 21-nt siRNAs, 24-nt siRNAs that accumulated in *ddm1 rdr2* were decreased in *ccdr* (Figure 3A), and 24-nt siRNAs produced from *ATHILA* TEs in *ddm1 rdr2* were severely reduced in *ccdr* (Figures 3B-3D). We confirmed that these CCR4-dependent 24-nt siRNAs were produced in an RDR6-dependent manner (Figures 3D-3F and S4B). In addition, 24-nt siRNA levels at *TAS* gene loci were also decreased in *ccr4* as observed in *rdr6* (Figure S7). Collectively, these results demonstrate that CCR4 plays a critical role in the RDR6-dependent siRNA biogenesis pathway.

**Figure 3.**
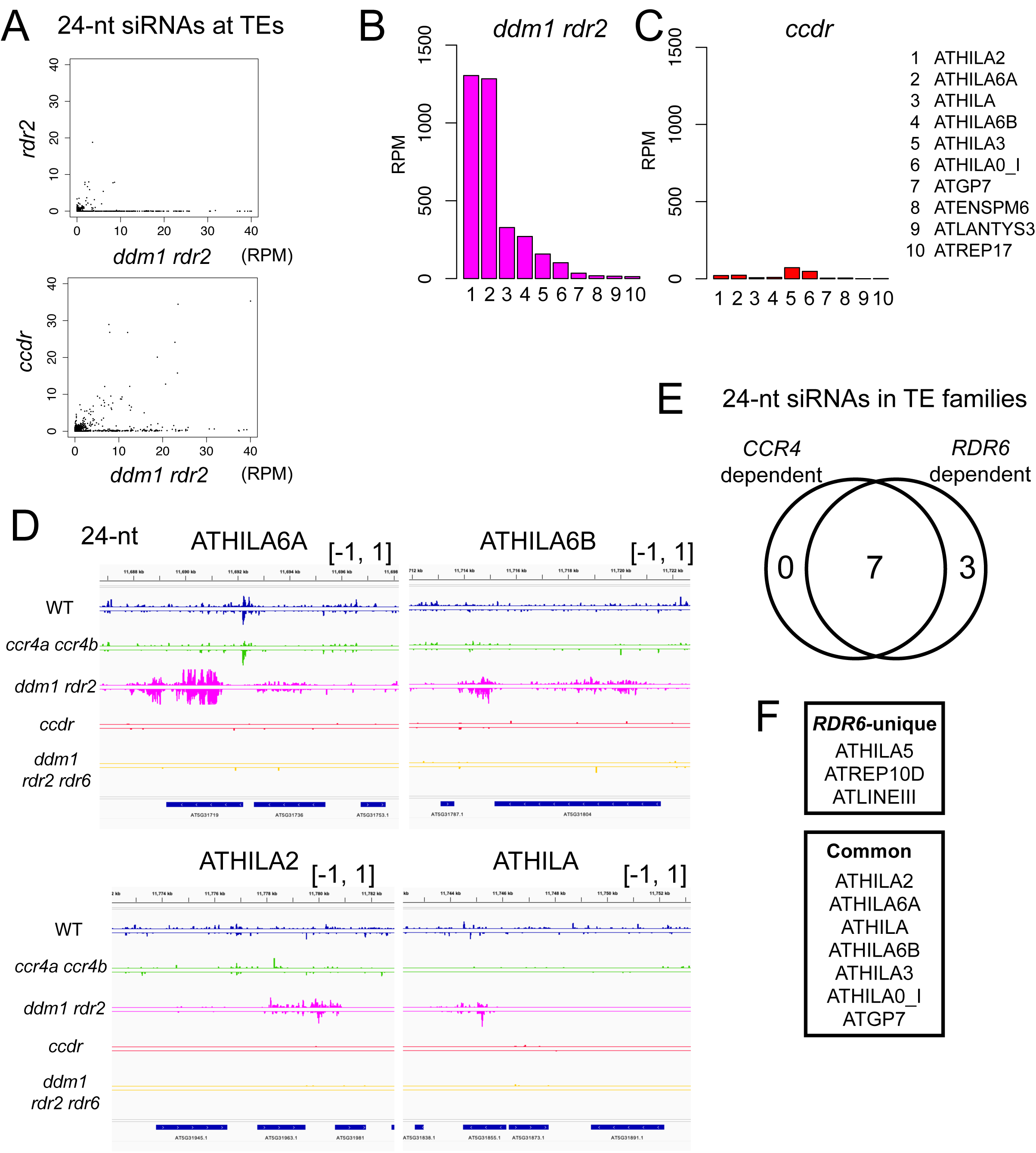
CCR4 acts in the RDR6-dependent 24-nt siRNA synthesis pathway. (A) Scatterplots comparing normalized 24-nt siRNA read counts (RPM) across *rdr2*, *ddm1 rdr2*, and *ccdr*. (B-C) The top 10 TE families associated with the highest levels (RPM) of 24-nt siRNAs in *ddm1 rdr2* (B), and the changes in 24-nt siRNA levels of these TEs in *ccdr* (C). (D) 24-nt siRNA levels (RPM) at *ATHILA* family retrotransposons. (E) Overlap between CCR4-dependent siRNAs and RDR6-dependent 24-nt siRNAs. TEs associated with 24-nt siRNAs (RPM > 20 in *ddm1 rdr2*)) were analyzed. The Venn diagram shows the number of TE families whose 24-nt siRNAs were reduced by more than half in *ccdr* or *ddm1 rdr2 rdr6* specifically, or in both mutants compared to *ddm1 rdr2*. (F) The list of TE families shown in Figure 3E.

### CCR4 shortens poly(A) tails of substrate RNAs for RDR6-dependent dsRNA synthesis

The above results suggest that, in the absence of CCR4, the single-stranded template RNAs for dsRNA synthesis become hyper-polyadenylated, and the presence of longer poly(A) tail inhibits efficient dsRNA synthesis by RDR6, as previously repored^23^. To investigate this further, we assessed the impact of *CCR4* loss on the poly(A) tail length of mRNAs from *ATHILA* TEs and *TAS* loci usnig Oxford Nanopore direct RNA sequencing (ONT-DRS). We first confirmed that the overall poly(A) tail length of mRNAs in *ccr4a ccr4b* is longer compared to those in WT (Figure 4A-4E), consistent with the previous report^18^. Importantly, the proportion of short poly(A)-tailed RNAs (with poly(A) tails less than 20 bp) is significantly smaller in the *ccr4a ccr4b* background (Figures 4A-4E), indicating that CCR4 is essential for generating short poly(A)-tailed mRNAs by trimming the poly(A) tail. Further analyses of poly(A) tail length of mRNAs from *ATHILA* TE loci showed that loss of *CCR4* led to significant reductions in short poly(A)-tailed *ATHILA2*, *ATHILA5* and *ATHILA6A/B* transcripts (Figures 4F-4H). Despite the low number of mapped reads in *TAS* loci, reductions in short poly(A)-tailed *TAS* transcripts were also observed in most *TAS* loci (Figure S8). These results suggest that CCR4 is crucial for the production of short poly(A)-tailed RNAs, which are the preferred substrates for RDR6-dependent dsRNA synthesis.

**Figure 4.**
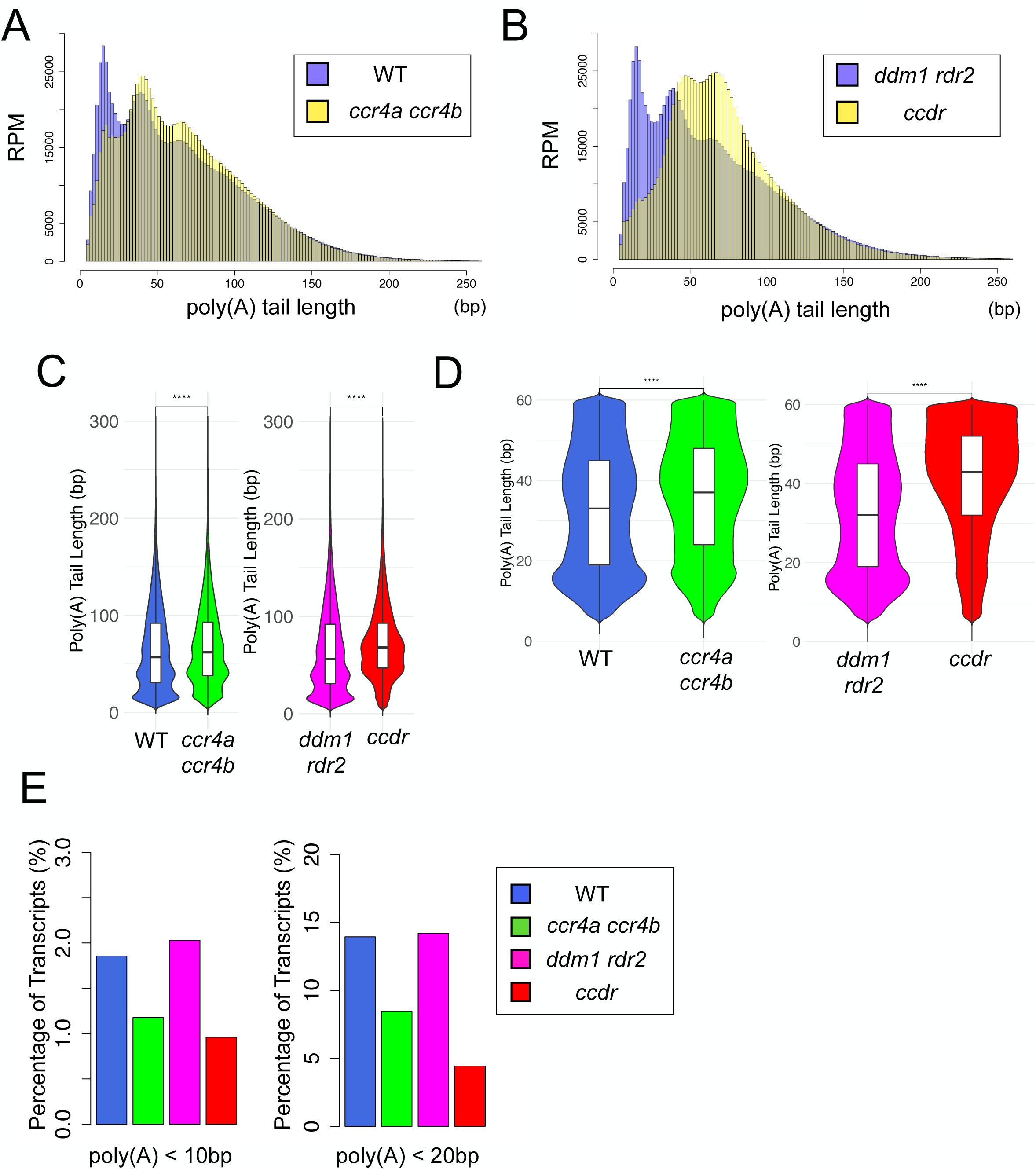
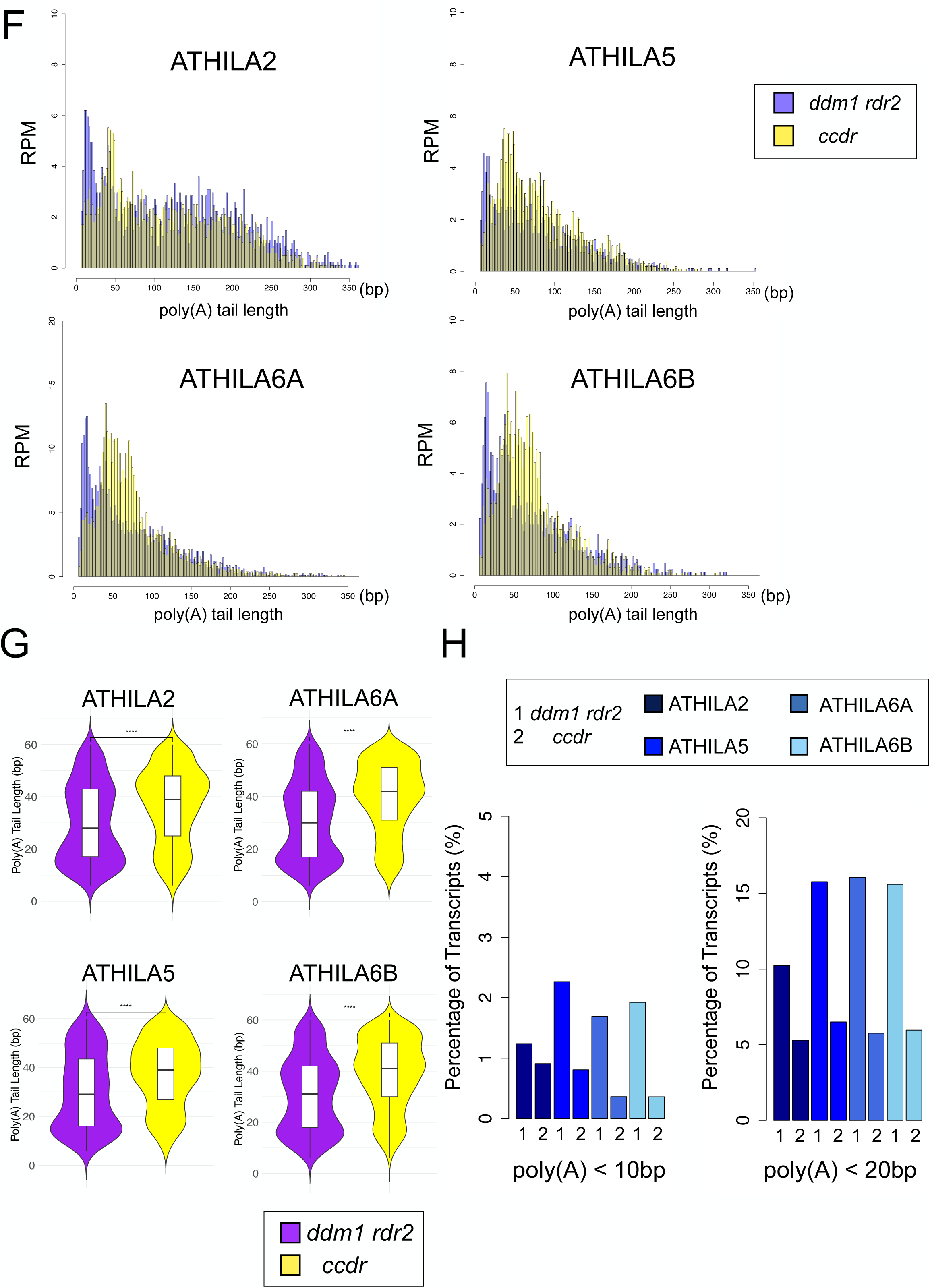
Poly(A) tail length measurement by Nanopore Direct RNA Sequencing. (A-B) Distribution patterns of poly(A) tail length of all transcripts in the indicated lines (WT and *ccr4a ccr4b* (A), *ddm1 rdr2* and *ccdr* (B)). Each bar in the histogram represents a 2 bp bin. (C-D) Violin plots showing poly(A) tail length distribution comparing WT (blue), *ccr4a ccr4b* (green), *ddm1 rdr2* (pink), and *ccdr* (red) within the 0-300 bp (C) and 0-60 bp (D) ranges (****: *p* < 0.0001, Wilcoxon Rank-Sum Test). (E) Proportion of short poly(A)-tailed transcripts in the indicated lines. The Y-axis indicates the percentage (%) of ONT-DRS reads with short poly(A) tails (< 20 bp (Left) or < 10 bp (Right)) relative to all mapped reads. (F) Poly(A) tail length analysis for *ATHILA* TE RNAs by ONT-DRS. Each bar in the histogram represents a 2 bp bin. Distribution patterns of poly(A) tail length were compared between *ddm1 rdr2* (purple) and *ccdr* (yellow). (G) Violin plots showing poly(A) tail length distribution in *ATHILA* TE RNAs. Distribution patterns were compared between *ddm1 rdr2* (purple) and *ccdr* (yellow) within the 0-60 bp range (****: *p* < 0.0001, *: *p* < 0.05, ns: not significant, Wilcoxon Rank-Sum Test). (H) Proportion of short poly(A)-tailed transcripts in *ATHILA* TEs in *ddm1 rdr2* (1) and *ccdr* (2). The Y-axis indicates the percentage (%) of ONT-DRS reads with short poly(A) tails (< 20 bp (E) or < 10 bp (F)) relative to all reads mapped to the indicated gene.

### CCR4 regulates biogenesis of centeromere-derived siRNAs and promotes DNA methylation at the centromere

Small RNA-seq data revealed that the global levels of 24-nt small RNA were also reduced in *ccr4* and *rdr6* (Figure S9A). The 24-nt siRNA levels were mildly decreased at TEs in *ccr4a ccr4b* and *rdr6*, with similar reduction patterns observed in both mutants (Figure S9B), suggesting that very weakly transcribed TE RNAs might be processed into siRNAs by CCR4 and RDR6. Notably, we also found that 24-nt siRNAs derived from the centromeric 178 bp repeats were reduced in *ccr4* and *rdr6* (Figure 5A). The loss of *ddm1 rdr2* led to a further reduction of siRNAs at the centromere repeats, which was exacerbated in the *ccr4* and *rdr6* background (Figure 5B). Based on previous studies on centromere repeat transcription in WT *Arabidopsis*^27^, we speculated that the centromere repeat-derived transcripts might be processed into 24-nt siRNAs via CCR4 and RDR6 activities, similar to tasiRNAs and easiRNAs. Indeed, we detected a certain number of RNA-seq reads mapped to the centromere repeat sequences in WT RNA samples (Table S3). Although our ONT-DRS analysis for WT and *ccr4a ccr4b* could not detect transcripts derived from centromere repeat sequences, likely due to insufficient read coverages, we could detect the reads containing centromere repeat sequences in *ddm1 rdr2* and *ccdr* (Figure 5C). Similar to *ATHILA* TE RNAs, the proportion of short poly(A)-tailed transcripts derived from centromere repeats was significantly reduced in *ccdr* compared to *ddm1 rdr2* (Figures 5C-5E), supporting the idea that CCR4 regulates the poly(A) tail of centromeric transcripts. Thus, CCR4 promotes the synthesis of centromeric 24-nt siRNAs.

**Figure 5.**
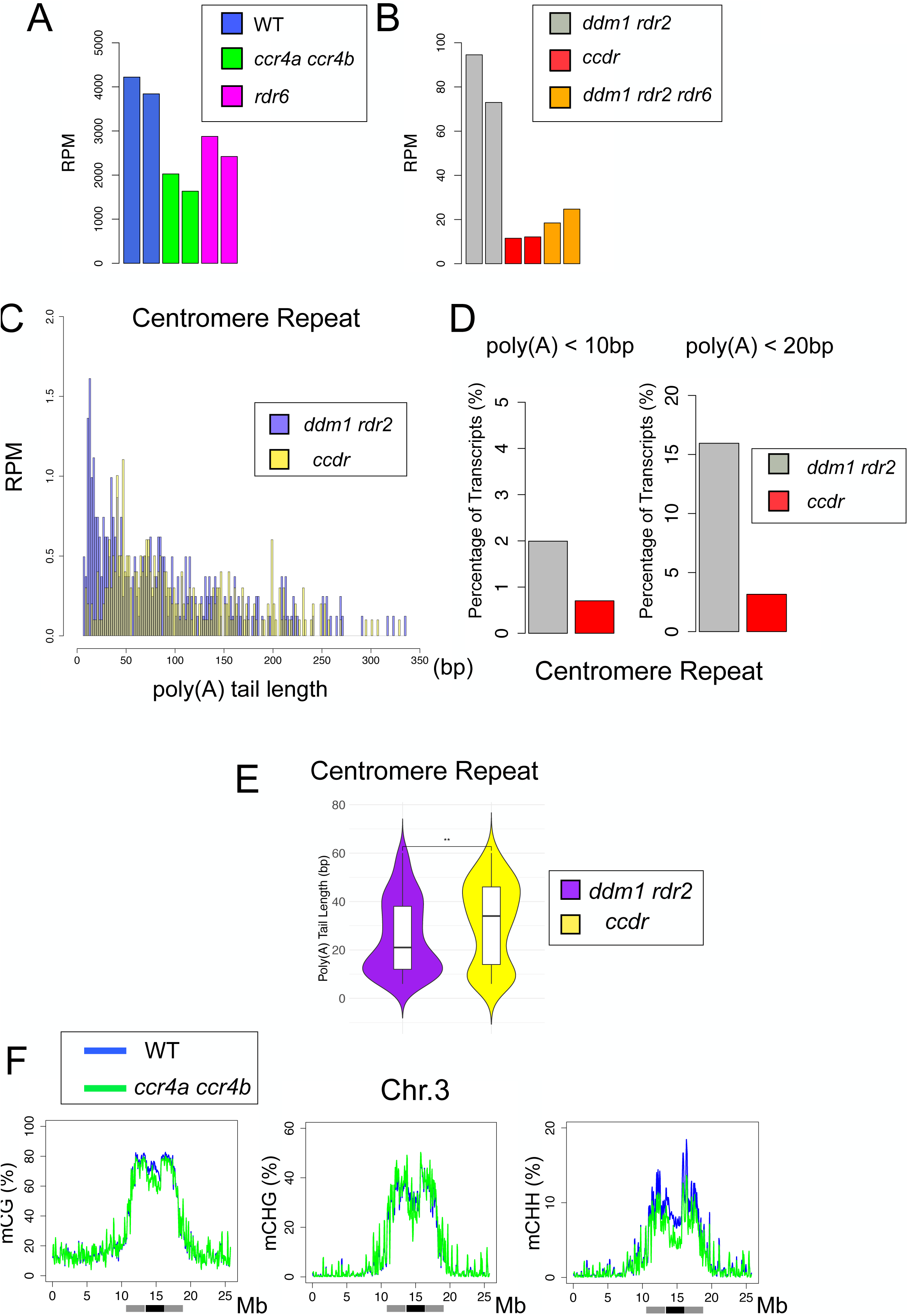
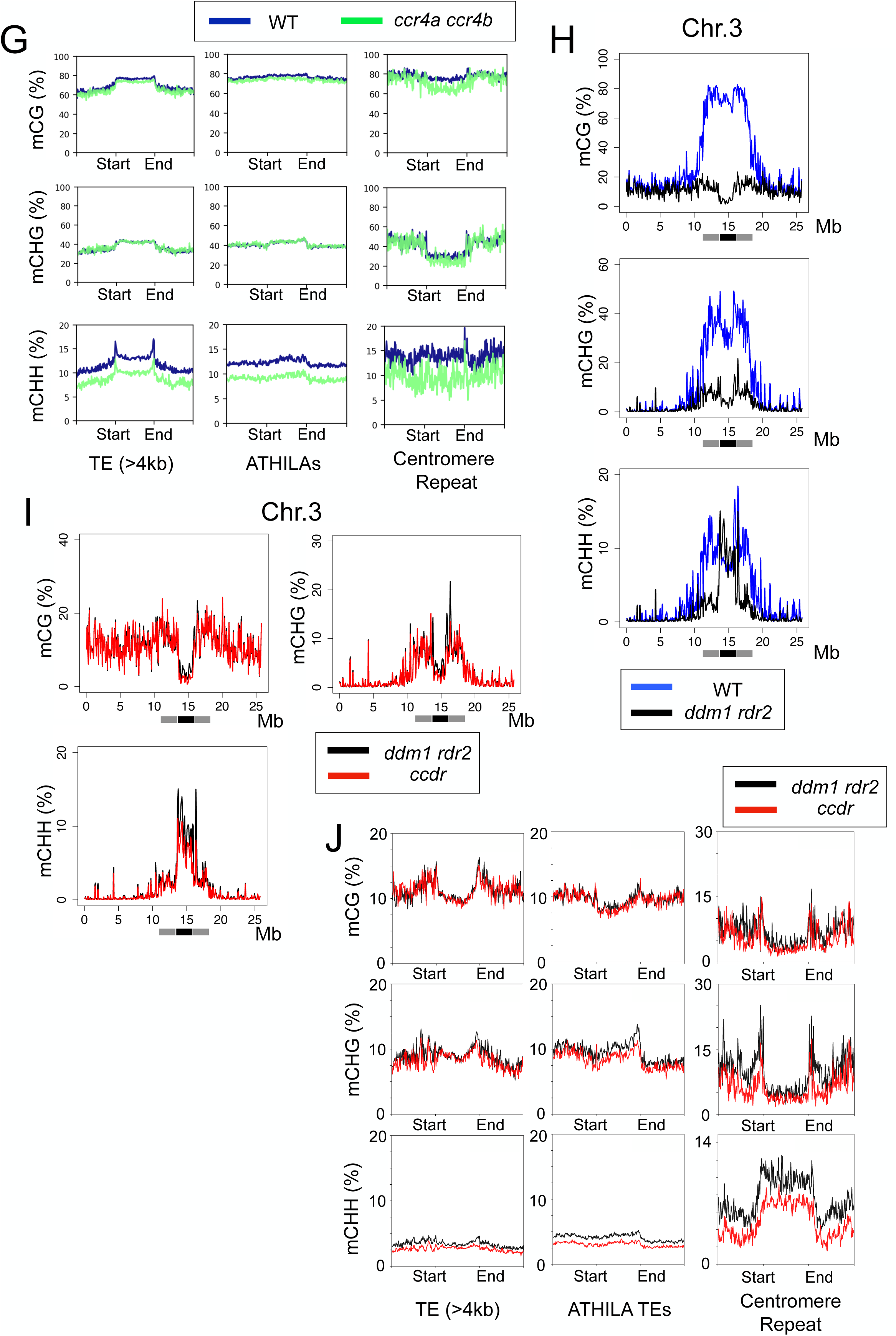
CCR4 promotes DNA methylation at the centromere repeats. (A) Normalized 24-nt siRNA levels (RPM) derived from centromere repeats detected in WT (blue), *ccr4a ccr4b* (green), and *rdr6* (pink). (B) Normalized 24-nt siRNA levels (RPM) derived from centromere repeats in *ddm1 rdr2* (gray), *ccdr* (red), and *ddm1 rdr2 rdr6* (orange). (C) Poly(A) tail length of centromere repeat-derived RNAs analyzed by ONT-DRS in *ddm1 rdr2* (purple) and *ccdr* (yellow). Each bar in the histogram represents a 2 bp bin. (D) Proportion of short poly(A)-tailed transcripts derived from centromere repeats in *ddm1 rdr2* (gray) and *ccdr* (red). The Y-axis indicates the percentage (%) of ONT-DRS reads with short poly(A) tails (< 20 bp (Left) or < 10 bp (Right)) relative to all mapped reads containing centromere repeat sequences. (E) Violin plots showing poly(A) tail length distribution in centromere repeat-derived transcripts. Distribution patterns were compared between *ddm1 rdr2* (purple) and *ccdr* (yellow) within the 0-60 bp range (****: *p* < 0.0001, **: *p* < 0.01, Wilcoxon Rank-Sum Test). (F) Metaplots of DNA methylation for mCG, mCHG, and mCHH in Chr. 3 of WT (blue) and *ccr4a ccr4b* (green). Gray and black bars indicate pericentromeric regions and centromere repeat domains, respectively. (G) Metaplots of DNA methylation (mCG, mCHG, and mCHH) over TEs (> 4 kb), *ATHILA* TEs, and centromere repeats in WT (blue) and *ccr4a ccr4b* (green). (H-I) Metaplots of DNA methylation for mCG, mCHG, and mCHH in Chr. 3 of WT (blue) and *ddm1 rdr2* (black) (H), and *ddm1 rdr2* (black) and *ccdr* (red) (I). Gray and black bars indicate pericentromeric regions and centromere repeat domains, respectively. (J) Metaplots of DNA methylation (mCG, mCHG, and mCHH) over TEs (> 4 kb), *ATHILA* TEs (> 4 kb), and centromere repeats in *ddm1 rdr2* (black) and *ccdr* (red).

Given that changes in 24-nt siRNA levels are likely to affect DNA methylation at non-CG contexts (mCHG and mCHH; H = C, A, or T), we next examined whether CCR4 regulates DNA methylation in the centromeric regions of chromsomes. Consistent with the decreased 24-nt siRNA levels, *ccr4* exhibited reduced mCHH levels in both the peri-and centromeric regions (Figures 5F and S10A). Specifically, mCHH levels were decreased at both TEs within the centromeres and centromere repeats in *ccr4* mutants (Figures 5G and S10B). Since *ccr4* exacerbated the developmental phenotypes of *ddm1 rdr2* and caused a further reduction of centromeric siRNAs (Figures 1 and 5B), we speculated that CCR4-dependent siRNAs maintain centromeric DNA methylation independently of *DDM1* and *RDR2*. Indeed, although *ddm1 rdr2* mutants exhibited globally reduced DNA methylation in both CG and non-CG contexts, mCHH levels remained mostly unaffected at the core centromeric regions (Figures 5H and S11). Importantly, we observed that mCHH methylation at the core centromeric regions was further reduced in *ccdr* compared to *ddm1 rdr2* (Figures 5I, J and S12). There was little difference in DNA methylation between *ddm1 rdr2* and *ccdr* at general TEs including *ATHILA* TEs (Figure 5J). However, *ATHILA2* and *ATHILA6 A/B*, which are associated with CCR4-dependent 24-nt siRNAs, showed lower mCHH levels in *ccdr* than in *ddm1 rdr2* (Figures 3B, 3C and S13A). We also found that loci associated with CCR4-dependent 24-nt siRNA generally lost non-CG DNA methylation in *ccdr* (Figures S13B-13E and Table S4). Based on these results, we concluded that CCR4-dependent 24-nt siRNAs maintain non-CG DNA methylation over centromeric repeats and TEs in the absence of the primary DNA methylation pathways.

### CCR4 safeguards centromeric heterochromatin and chromosome stability

Given that DNA methylation recruits histone H3K9 methyltransferases to chromatin in *Arabidopsis*^28^, we next examined the impact of *ccr4* on histone H3K9 dimethylation (H3K9me2) by ChIP-seq. H3K9me2 is a heterochromatic histone modification critical for centromere function in other organisms^29–31^. We observed that H3K9me2 is highly enriched at the centromeres in WT, but was reduced at the pericentromere in *ddm1 rdr2* (Figures 6A and 6B), while the mutant retained high levels of H3K9me2 at the centromere core (Figures 6B and S14), consistent with the retention of mCHH in that region (Figure 5H). The *ccr4* mutation slightly decreased centromeric H3K9me2 levels, and it caused a further reduction in H3K9me2 in the absence of DDM1 and RDR2 in *ccdr* (Figures 6A-6C and S14). The affected regions largely corresponded to centromere repeats, where mCHH levels were further diminished in *ccdr* compared to *ddm1 rdr2* (Figures 6C and 6D). Although mCHH levels are generally lower than mCG and mCHG in the *Arabidopsis* genome (Figures 5F and S10A), the frequency of CHH sites in the genome is much higher than that of CG and CHG sites (Figure S15). Centromere repeats have a higher frequency of CHH sites compared to the average in the whole genome or in *ATHILA* TE sequences, while less CG and CHG sites (Figure S15). This may explain the larger impact of mCHH on H3K9me2 at the centromere repeats. Centromeric H3K9me2 levels in *ddm1 rdr2 rdr6* were also decreased compared to *ddm1 rdr2*, but the reduction is less significant than in *ccdr* (Figure S16), suggesting that CCR4 may promote centromeric H3K9 methylation in both an RDR6-dependent and independent manner. H3K9 methylation plays important roles in centromeric chromatin condensation and mitotic chromosome stability^32, 33^. Indeed, we observed a disturbed centromere structure in *ccdr* nuclei, where *ccdr* exhibited severely misaligned chromosomes during metaphase (Figure 6E). In contrast, mutants defective in either CCR4 or DNA methylation pathways showed minor defects, consistent with the high levels of H3K9me2 retained at the centromere core (Figure 6A). The severe chromosome instability in *ccdr* may explain the aggravated developmental defects in the mutant. Taken together, CCR4 and primary DNA methylation pathways cooperatively maintain centromeric H3K9 methylation, which is essential for mitotic chromosome stability and development.

**Figure 6.**
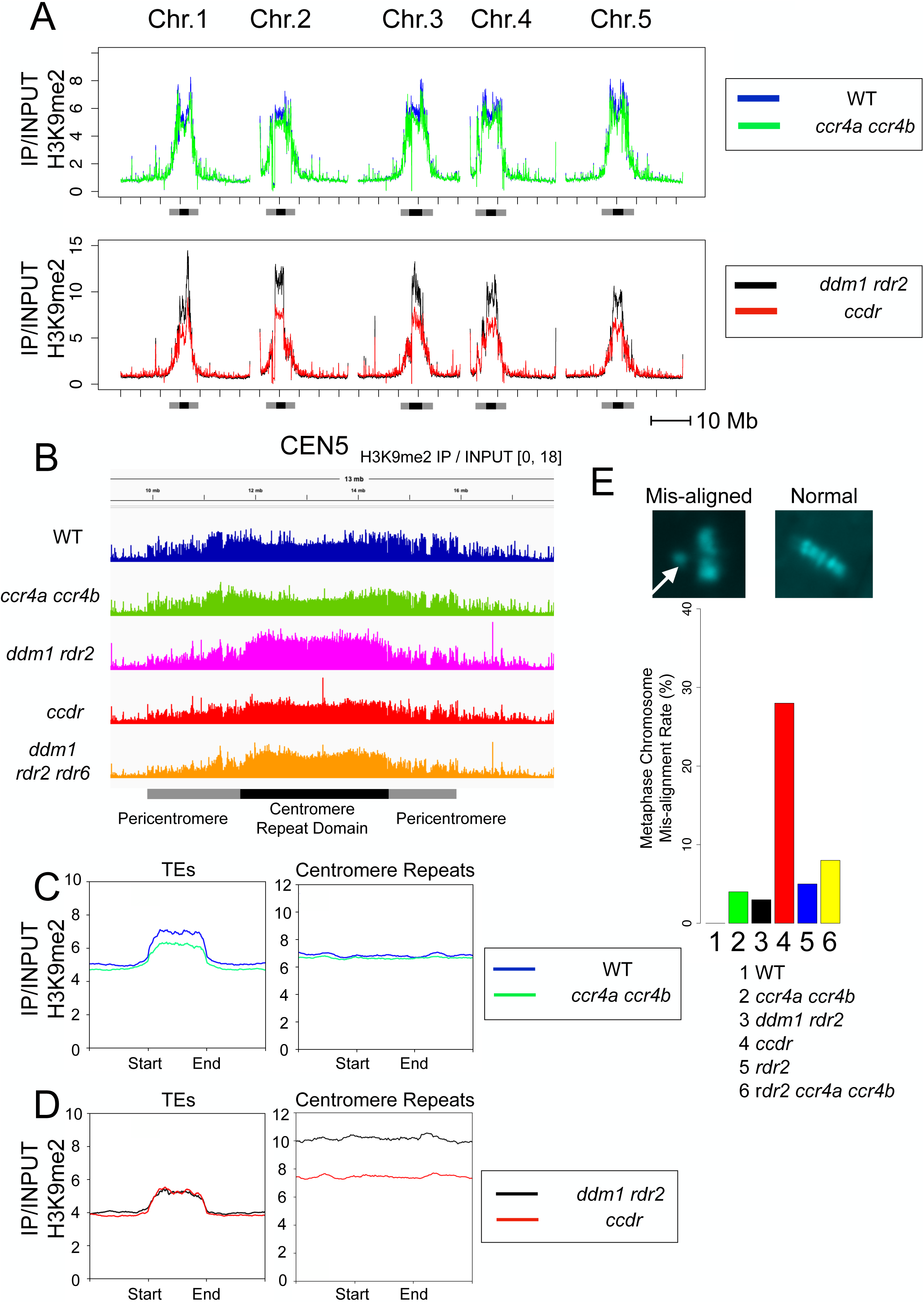
CCR4 and DNA methylation maintain centromeric H3K9me2 and chromosome stability. (A) Genome-wide H3K9me2 levels compared between WT (blue) and *ccr4a ccr4b* (green) (upper panel), and *ddm1 rdr2* (black) and *ccdr* (red) (lower panel). H3K9me2 ChIP-seq signals were normalized to input signals (IP/INPUT). Gray and black bars indicate pericentromeric regions and centromere repeat domains, respectively. (B) Normalized H3K9me2 signal (IP/INPUT) tracks over the chromosome 5 centromere in the indicated lines. (C-D) Metaplots of H3K9me2 signals (IP/INPUT) over TEs (> 4 kb) and centromere repeats in the indicated lines. (E) Percentage of misaligned metaphase chromosomes in the indicated mutants. One hundred metaphase cells were analyzed for each line. Inset images show representative misaligned chromosomes (white arrow) and normal chromosomes stained with DAPI.

## Discussion

In this study, we demonstrated the crucial role of CCR4 in poly(A) shortening for siRNA synthesis and centromere integrity in *Arabidopsis*. Our findings of the involvement of CCR4 in the RDR6-dependent siRNA biogenesis align with previous reports that RDR6 preferentially uses short poly(A)-tailed or poly(A)-less template RNA for dsRNA synthesis *in vitro*^23^. This suggests that plants differentiate between deadenylated RNAs targeted by RNAi and functional polyadenylated mRNAs.We revealed that CCR4 maintains centromeric DNA methylation and H3K9 methylation independently of the primary DNA methylation pathways by promoting RDR6-dependent siRNA synthesis. Although pericentromeric heterochromatin is diminished in *ddm1 rdr2* (Figure 6B and S14), the mutant retains H3K9 methylation and mCHH over the centromere repeats, likely contributing to the preservation of functional centromeres and preventing severe chromosome instability. A recent study demonstrated that RDR6-dependent easiRNAs originating from the *ATHILA5* TE promote the mitotic segregation of the *Arabidopsis* chromosome 5^20^. While we did not specifically investigate whether the chromosome instability and developmental phenotypes in *ccdr* result from *ATHILA5*, it is noteworthy that *ccdr* retains relatively more *ATHILA5*-derived siRNAs than *ddm1 rdr2 rdr6* (Figure 3E, F and S3B). Thus, the developmental and chromosomal defects in *ccdr* may be primarily attributed to the globally reduced centromeric siRNAs, mCHH and H3K9me2 across all the chromosomes.

Our data indicated that *ccr4* and *rdr6* cause similar impacts on small RNAs, epigenomes and phenotypes, but we also observed that the effect of *ccr4* differs in part from those of *rdr6*. In the absence of DDM1 and RDR2, *ccr4* caused a greater reduction of H3K9 methylation on the centromere repeats compared to *rdr6* (Figure S16). This observation is reminiscent of the previous study in yeast, which demonstrated that the *S. pombe* CCR4-NOT complex degrades chromatin-bound centromeric RNAs to facilitate access for H3K9 methyltransferase to the chromatin^20^. Importantly, this pathway acts independently of RNAi^19^. Our result suggests that, similar to *S. pombe*, *Arabidopsis* CCR4 may also process chromatin-bound RNAs on the centromere repeat to promote H3K9 methylation in an RNAi-independent manner.

Our study provided the first evidence that the CCR4-NOT complex promotes chromosome stability in plants. There is also growing evidence in other organisms that CCR4-NOT plays a role in controlling chromosome stability. In *S. pombe*, CCR4-NOT mutants exhibit reduced viability in aneuploid cells^34^. In *S. cerevisiae*, CCR4-NOT is implicated in maintaining rDNA copy number by degrading non-coding RNA on the rDNA chromatin^35^, and similar instability of rDNA has been reported in *S, pombe*^18^. In human, depletion of CCR4-NOT components causes transcription-induced genome instability^36^. Hence, RNA quality control is essential not only for regulation of gene expression but also for maintenance of chromosome integrity.

While DNA methylation has been shown to maintain centromeric heterochromatin, there has been limited evidence om how plant centromere function is maintained, due to a lack of studies directly demonstrating the effect of DNA methylation on chromosome stability. Our study sheds light on the elaborate mechanisms that safeguard centromere function through multiple epigenetic layers, which are essential for chromosome integrity, cell proliferation and plant development.

## Methods

### Plant Materials and growing condition

All the analyzed *Arabidopsis thaliana* lines are Columbia-0 (Col-0) accession, and plants were grown under long-day conditions (light 16h, dark 8h) at 22C. For knockout mutations, the following T-DNA lines and point mutants were used: *ccr4a* (SAIL1231_B04), *rdr2* (SALK_059661), *ddm1-1*^37^, *rdr6-11*^38^.

### Short read sequencing

Short read sequencing was performed for ChIP-seq, RNA-seq, small RNA-seq and Enzymatic Methyl-seq on an Illumina NovaSeq6000 platform with 150 bp paired-end reads in OIST Sequencing Section.

### ChIP-seq

ChIP-seq was performed with two biological replicates using the protocol used in the previous study with some modifications^20, 39^. 0.1 g of frozen 12-day-old seedlings were ground under liquid nitrogen, and the ground tissues were cross-linked with 12.5 ml of formaldehyde solution (1% Formaldehyde, 10 mM HEPES pH 7.6, 1M sucrose, 5 mM KCl, 5 mM MgCl_2_, 5 mM EDTA pH 8.0, 0.6% Triton-X100, 0.1% 2-mercaptoethanol, 1x complete protease inhibitor (Sigma)) for 10 minutes at room temperature. Cross-linking reaction was quenched by adding 0.85 ml of 2 M glycine and samples were incubated for 5 minutes at room temperature. The ground tissues were collected by centrifuging at 3000 x g for 10 min at 4 C followed by resuspension in 200 µl of buffer 1 (50 mM HEPES-KOH (pH 7.6), 140 mM NaCl, 1 mM EDTA, 1% Triton X-100, 0.1% Na-Deoxycholate, 1x complete protease inhibitor) containing 1% SDS. The samples were incubated at 4C for 10 minutes and diluted with 2 ml of buffer 1. Chromatin was sheared to 250-500 bp with Branson Digital Sonifier (30 sec ON and 59.9 sec OFF for 12 cycles). After the centrifuge at 15,000 rpm and 4C for 10 minutes, the supernatant was used for immuno-precipitation. Dynabeads M-280 Sheep anti-Mouse IgG (11202D, Thermofisher Scientific) and the antibody against H3K9me2 (MABI0307, MAB Institute, Inc.) were used for immuno-precipitation. Immunoprecipitates were washed twice with 0.5 ml of buffer 1, twice with 0.5 ml of buffer 1’ (50 mM HEPES-KOH (pH 7.5), 500 mM NaCl, 1 mM EDTA, 1% Triton-X100, Na-Deoxycholate), once with 1 ml of buffer 2 (10 mM Tris-HCl (pH 8.0), 250 mM LiCl, 0.5% NP-40, 0.5% Na-Deoxycholate), and once with 1 ml of TE buffer. Input and washed immunoprecipitated samples resuspended in 100 µl of TE buffer were treated with 0.1 mg/ml RNase A at 37 C for 30 minutes and with 0.25 mg/ml Proteinase K and 0.25% SDS at 42 C for 1hr. Samples were then reverse-crosslinked at 65 C overnight, followed by purification with QIAquick PCR purification kit (QIAGEN 28106). Library preparation was performed using NEB Next Ultra II DNA Library Prep Kit (E7645) and NEBNext® Multiplex Oligos for Illumina (E6440S), following the manufacturer’s instructions. For sequencing data analysis, sickle (v1.33) was used to check the quality of sequencing reads, and the filtered reads were aligned to the *Arabidopsis* reference genome (TAIR10 and ColCEN T2T assembly) using bowtie2 (v2.4.2)^40^. PCR duplicates were removed with Samtools^41^, and the processed bigwig and bedGraph files normalized with RPM were generated with deepTools (v3.4.3) and bedtools (v2.29.2), respectively.

### RNA-seq

Total RNA was extracted from the 12-day old plant seedlings using Direct-zol RNA Miniprep kit (Zymoresearch, R2050). Library preparation was performed using NEBNext Ultra II Directional RNA Library Prep Kit (NEB, E7760) and QIAseq FastSelect rRNA Plant Kit (334317) following manufacturer’s instructions. For the sequencing data analysis, sickle (v1.33) was used to check the quality of sequencing reads, and the filtered reads were aligned to the *Arabidopsis* reference genome (TAIR10 and ColCEN T2T assembly) obtained from TAIR (https://www.arabidopsis.org) using STAR (v2.7.9a)^42^. Normalized processed files were generated with deepTools (v3.4.3)^43^. Differentially expressed gene analysis was performed among three biological replicates of WT and mutants using Exact Tests in edgeR (v3.36.0) (FDR<0.01)^44^.

### Oxford Nanopore direct RNA sequencing

Total RNA was extracted from the 12-day old plant seedlings using Direct-zol RNA Miniprep kit (Zymo research, R2050). Library preparation was performed using Nanopore Direct RNA Sequencing Kit (SQK-RNA004) followed by sequencing with PromethION RNA flow cell (FLO-PRO004RA). For data analysis, Dorado (v0.5.3) was used for basecalling and poly(A) tail estimation. Minimap2 (v2.28-r1209) was used for mapping reads to the *Arabidopsis* ColCEN T2T genome assembly^45^. Reads mapped to specific genes were extracted with Samtools.

### Small RNA-seq

Total RNA was extracted from 5-week-old plant leaves using Direct-zol RNA Miniprep kit (Zymo research, R2050). Library preparation was performed using NEXTFLEX® Small RNA-Seq Kit v3 (Cosmobio, 5132-06). For sequencing data analysis, removal of the adaptor sequence and read quality check were performed with Cutadapt and fastx trimmer implemented in FASTX_Toolkit (v0.0.13) (http://hannonlab.cshl.edu/fastx_toolkit/index.html)^46^. The filtered reads were then aligned to the *Arabidopsis* reference genome (TAIR10 and ColCEN T2T assembly) with Bowtie2, and the processed bedGraph files normalized with RPM were generated with Samtools and bedtools (v2.29.2)^47^. Read counts for each gene and TE were obtained using featureCounts (v1.5.2) with default settings, and two biological replicates were analyzed to confirm reproducibility of the results^48^.

### Enzymatic methyl-seq

Genomic DNA was extracted from 6-week-old plant leaves using Nucleon phytopure (Sigma-Aldrich, RPN8510), and the extracted DNA was sheared into 400 bp with Covaris M220. NEBNext Enzymatic Methyl-seq Kit (NEB, E7120L) and Multiplex Oligos for Enzymatic Methyl-seq (E7140S) were used for library preparation. For sequencing data analysis, sickle (v1.33) was used to check the quality of sequencing reads, and the filtered reads were aligned to the *Arabidopsis* reference genome (TAIR10 and ColCEN T2T assembly) using Bismark (v0.22.3)^49^. Information of cytosine methylation was obtained by methylation extractor implemented in the Bismark program.

### Microscope analysis

10-day old seedlings were treated with 1ug/ml DAPI and 0.1% Triton X-100, and the stained root-tip nuclei were analyzed with CARL ZEISS LSM780 confocal microscope. Flower photos were taken with Leica KL200 microscope.

### Plasmid construction for *CCR4B* knockdown

A partial fragment of the *CCR4B* gene and its inverted form separated with *GUS* spacer were amplified by PCR, followed by cloning into pPZP2H-p35S^20^. The plasmid was transformed into Agrobacterium tumefaciens (GV3101), and then introduced into *Arabidopsis thaliana* by the floral dip transformation method^50^. Primers used to generate the plasmid are listed in Table S5.

## Materials availability

Newly generated materials are made available upon request by contacting the lead contact.

## Data and Code Availability

The sequencing data used in this study have been deposited in the Gene Expression Omnibus (GEO) with the following accession codes: GSE274190 (ChIP-seq), GSE274191 (RNA-seq), GSE274192 (smallRNA-seq), GSE274196 (EM-seq), and GSE274200 (ONT-DRS). This study does not report the original code. Any additional information required to reanalyze the data reported in this paper is available from the lead contact upon request.

## Lead contact

Further information and requests for resources and reagents should be directed to and will be fulfilled by the lead contact, Atsushi Shimada

## Acknowledgements

This work was supported by MEXT Grant-in-Aid for Transformative Research Areas (A) JP20H05913 to H. S. and by OIST. We thank OIST Sequencing Section for ONT-DRS and Illumina sequencing services, Dr. Martienssen for sharing mutant seeds, and OIST IT Section for technical support in building the web interface to access data.

## Author Contributions

A. S. designed the study. A. S. performed experiments and data analysis. A. S. and H. S. prepared the manuscript.

## Supplemental information

Figures S1-S16 and Tables S1-S5

## Graphical abstract

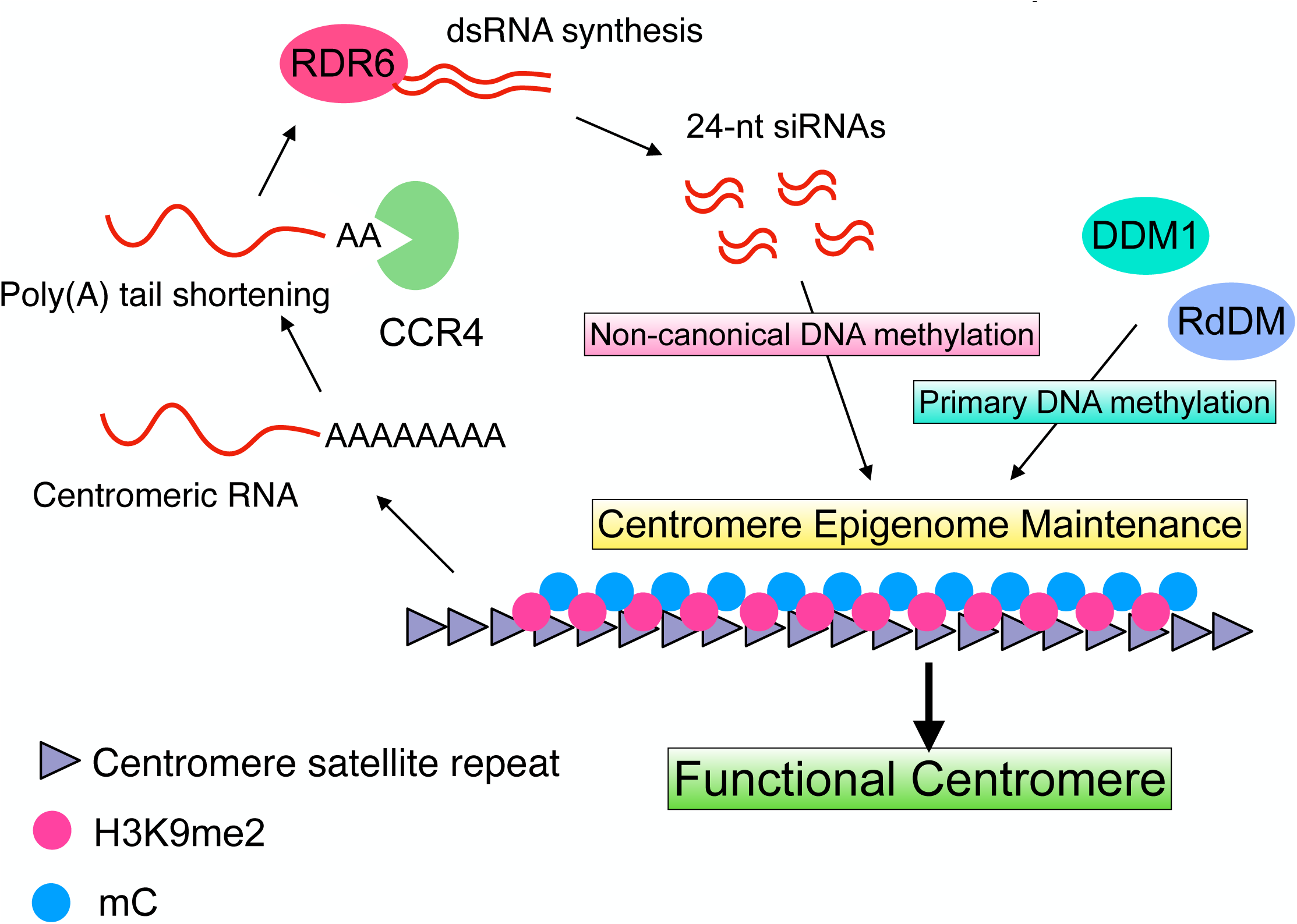

