## Supplementary Materials for "RNA quality control by CCR4 safeguards chromatin integrity and centromere function in *Arabidopsis*"

Figure S1

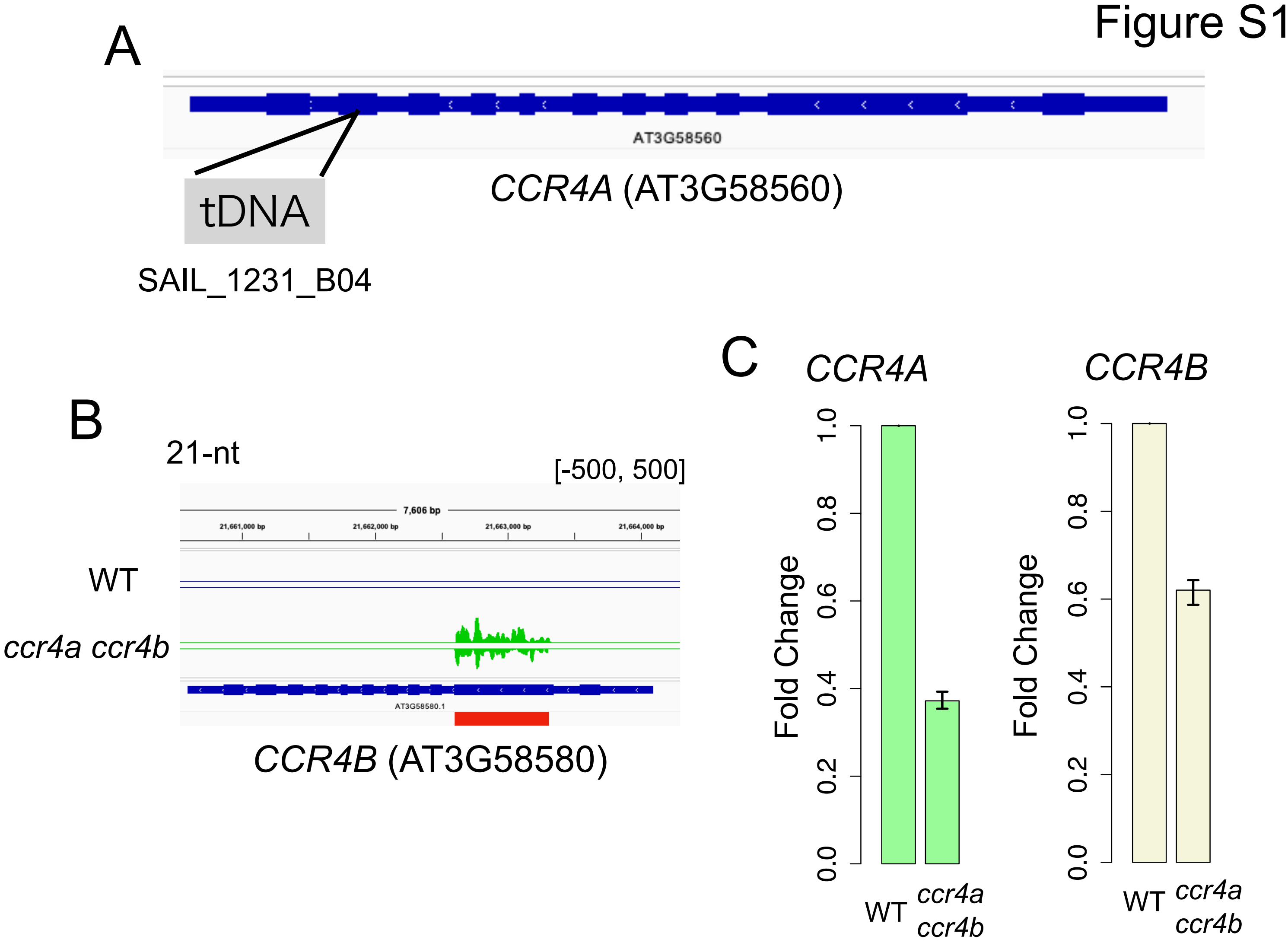


**Figure S1. Construction of *ccr4a ccr4b* mutant, related to Figure 1.**
(A) The T-DNA insertion site of *CCR4A* in SAIL_1231_B04 line.
(B) Tracks showing 21-nt small RNA derived from the hairpin RNA construct targeted for *CCR4B* knock-down in the *ccr4a* mutant background.
(C) Expression levels of *CCR4A* (Left) and *CCR4B* (Right) comparing between WT and *ccr4a ccr4b*. Y-axis indicates the ratio of normalized RNA-seq read count (CPM) in *ccr4a ccr4b* to that in WT (WT = 1). Three biological replicates were analyzed (Error bar = SEM, n=3).

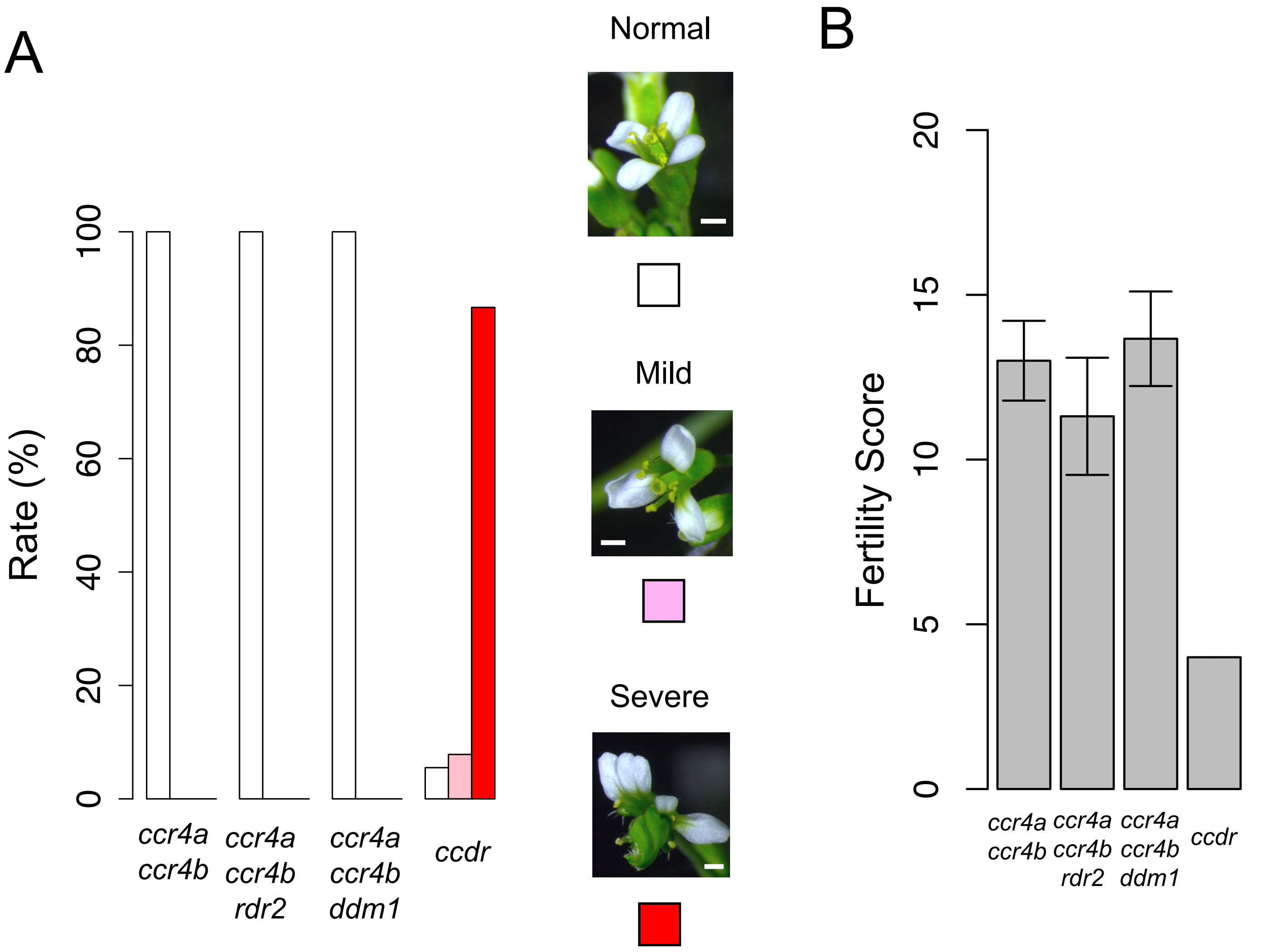


**Figure S2. Phenotypes of mutants defective in *CCR4* and DNA methylation**
(A-B) Flower phenotype (A) and fertility (B) in the indicated lines. Flowers were divided into three categories: Normal (white), Mildly defective (Mild; pink) and Severely defective (Severe; red), and 600 flowers from twelve T1 transformants (50 flowers/plant) were analyzed for each line. The white scale bar in the flower photos is 0.5 mm. Fertility Score represents the primary silique length (mm) with decimal numbers rounded down. T1 transformants were analyzed for *ccr4a ccr4b* (n = 16), *ccr4a ccr4b rdr2* (n = 16), *ccr4a ccr4b ddm1* (n = 12) and *ccdr* (n = 12) (Error bar = SD).

### Figure S3

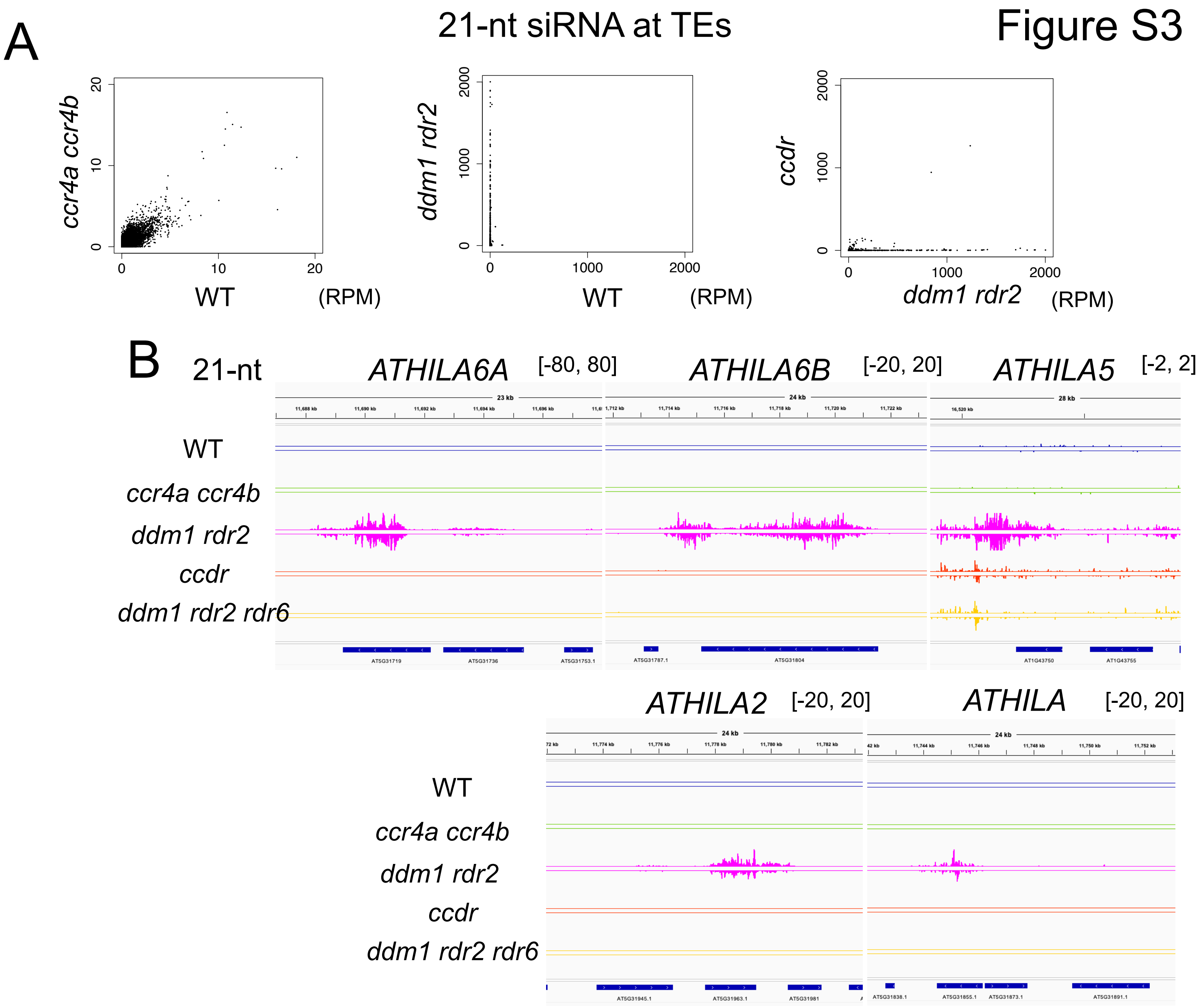
A
21-nt siRNA at TEs
ccr4a ccr4b
WT
(RPM)
ddm1 rdr2
WT
(RPM)
ccdr
ddm1 rdr2
(RPM)
B
21-nt
ATHILA6A [-80, 80]
ATHILA6B [-20, 20]
ATHILA5 [-2, 2]
WT
ccr4a ccr4b
ddm1 rdr2
ccdr
ddm1 rdr2 rdr6
ATHILA2 [-20, 20]
ATHILA [-20, 20]
WT
ccr4a ccr4b
ddm1 rdr2
ccdr
ddm1 rdr2 rdr6

Figure S3

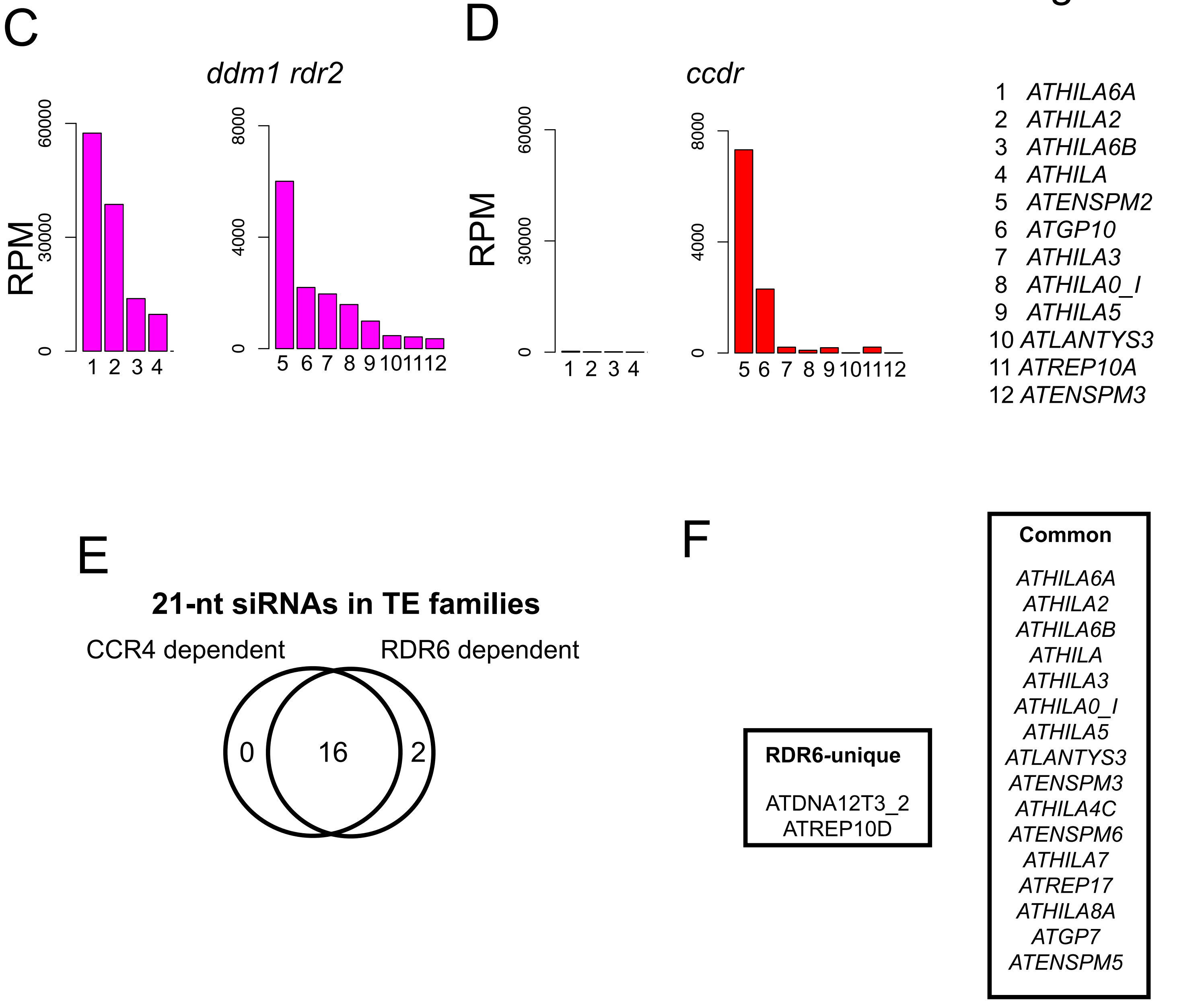


**Figure S3. CCR4 is essential for RDR6-dependent 21-nt easiRNA synthesis, related Figure 3.**
(A) Scatterplots comparing normalized 21-nt siRNA levels (RPM) from TEs across the indicated lines.
(B) 21-nt siRNA levels (RPM) at *ATHILA* retrotransposons in the indicated lines.
(C-D) The top 12 TE families that are associated with highest levels of 21-nt siRNAs in *ddm1 rdr2* (C), and the changes in 21-nt siRNA levels of these TEs were examined in *ccdr* (D).
(E) Overlaps between CCR4-dependent easiRNA-associated TE families and RDR6-dependent easiRNA-associated TEs. TEs associated with 21-nt easiRNAs (RPM > 20 in *ddm1 rdr2*)) were analyzed. The Venn diagram shows the number of TE families whose 21-nt siRNAs were reduced by less than half in *ccdr* or *ddm1 rdr2 rdr6* specifically, or in both mutants compared to *ddm1 rdr2*.
(F) The names of TE family in each category shown in Figure S3E.

Figure S4

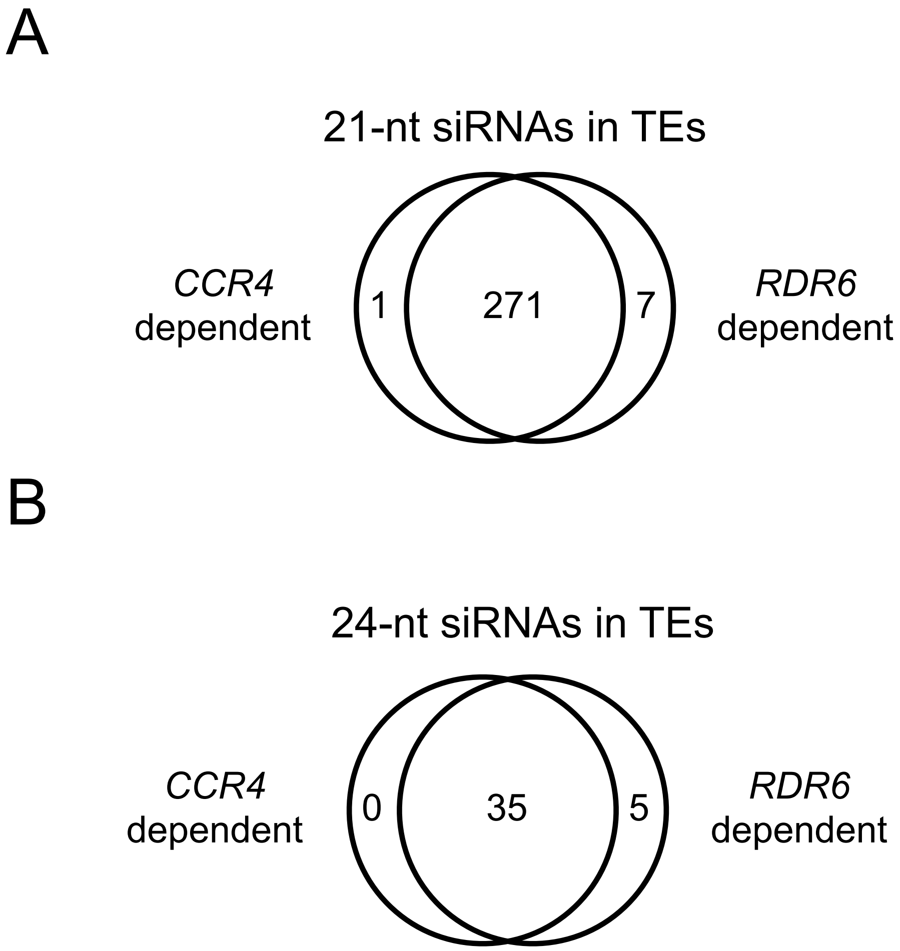


**Figure S4. Comparison of TE copies associated with CCR4-dependent siRNAs or RDR6-dependent siRNAs, related Figure 3.**
(A-B) Venn diagrams showing the number of TE copies whose siRNA synthesis is regulated by CCR4 or RDR6. TE copies associated with 21-nt (A) and 24-nt (B) siRNAs (RPM > 20 in *ddm1 rdr2*), showing a reduction to less than half of those in *ccdr* (or *ddm1 rdr2 rdr6*), are categorized as CCR4 (or RDR6)-dependent siRNA-producing TE copies.

### Figure S5

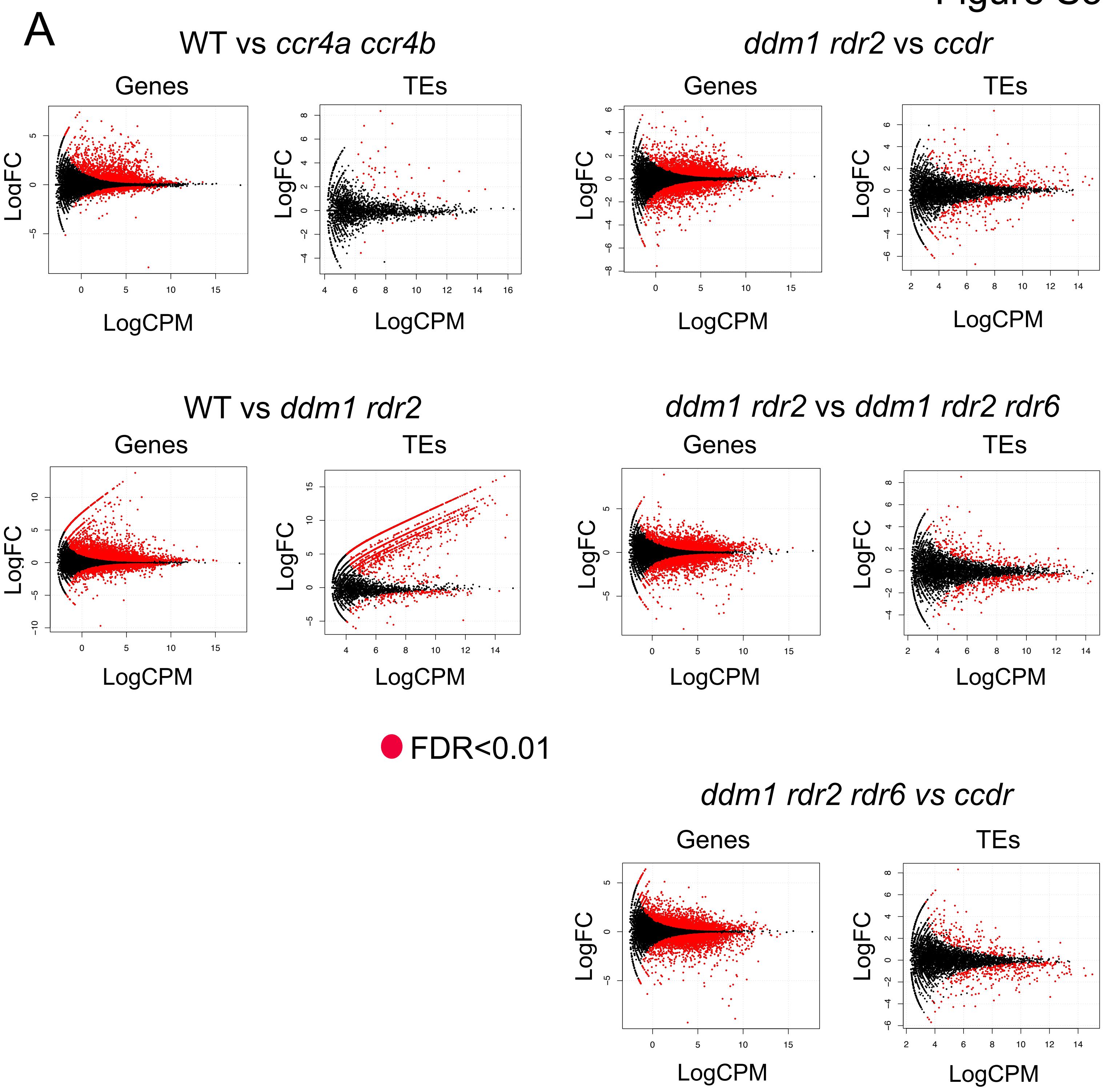
A
WT vs *ccr4a ccr4b*
*ddm1 rdr2* vs *ccdr*
WT vs *ddm1 rdr2*
*ddm1 rdr2* vs *ddm1 rdr2 rdr6*
*ddm1 rdr2 rdr6 vs ccdr*
Genes
TEs
LogFC
LogCPM
FDR<0.01

Figure S5

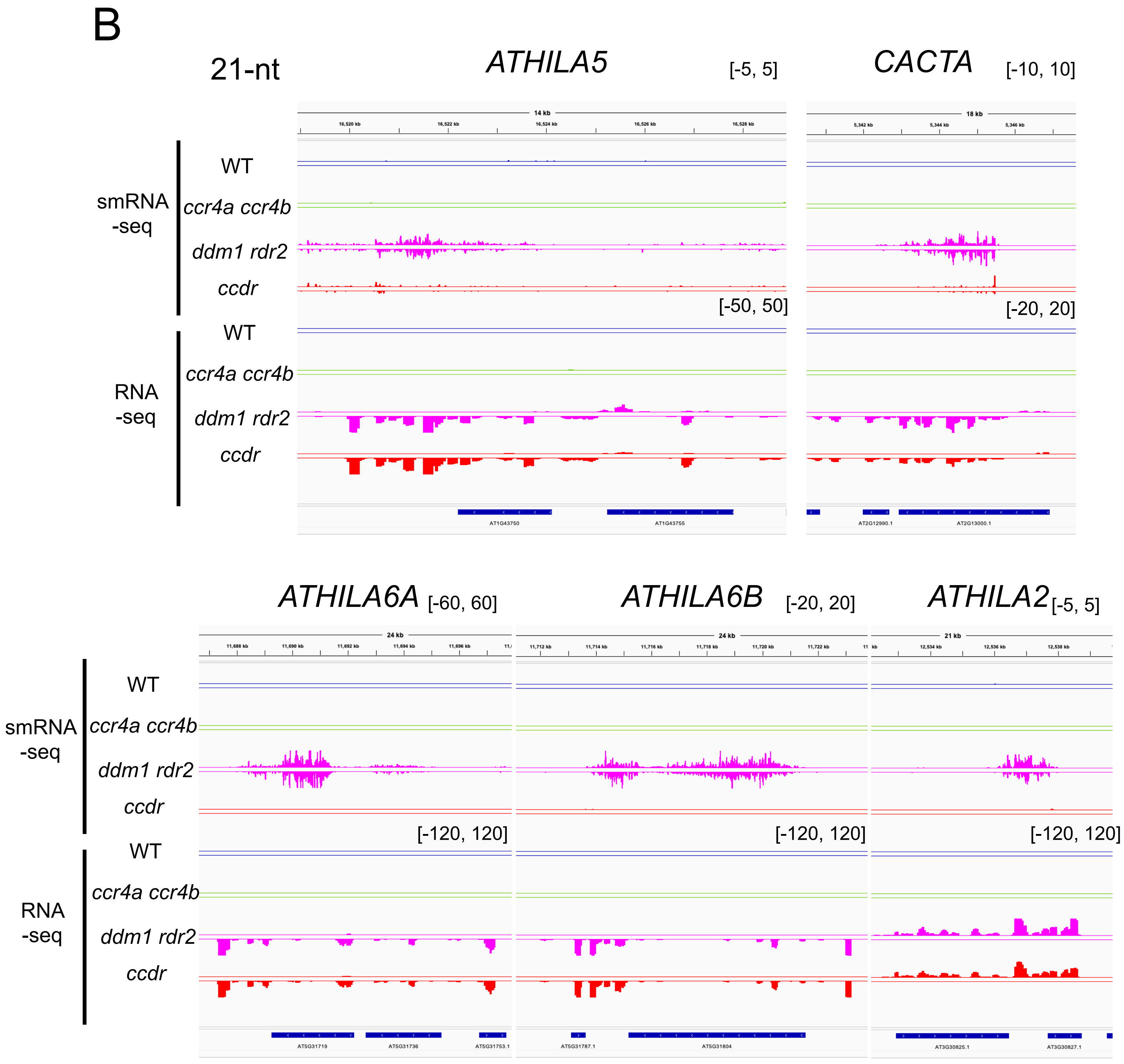


**Figure S5. Effect of *CCR4* mutation on gene and TE expression patterns.**
(A) Volcano plots analysis for differentially expressed genes, comparing across the indicated lines. Genes and TEs with significant difference (FDR < 0.01) are shown with red-colored dots. Three biological replicates of RNA-seq data were analyzed.
(B) RNA-seq and 21-nt small RNA-seq tracks over *ATHILA* and *CACTA* TEs in the indicated lines.

### Figure S6

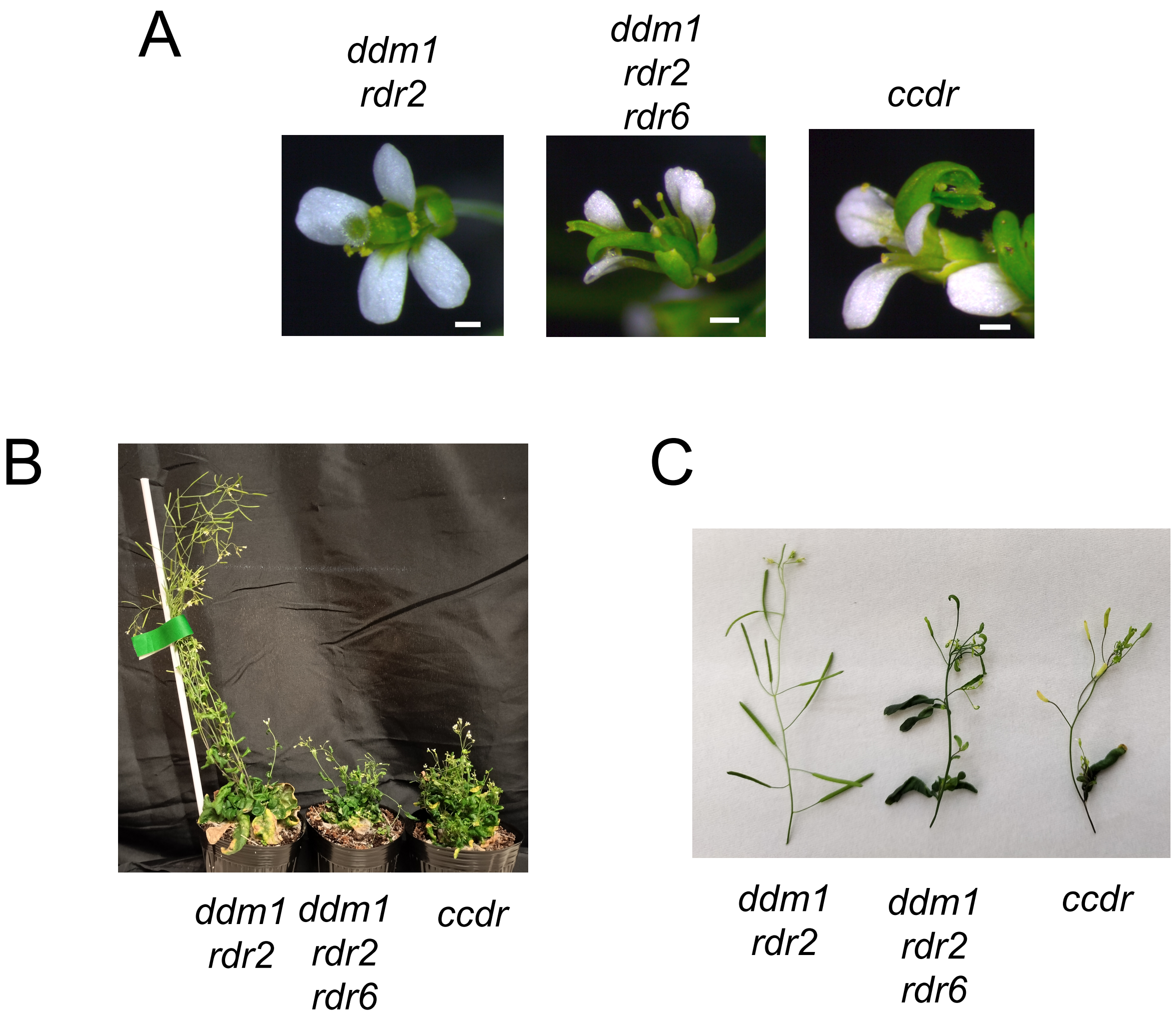


**Figure S6. Similarity of developmental phenotypes between *ccdr* and *ddm1 rdr2 rdr6*, related to Figure 3.**
(A) Flower photos of the indicated mutants. The white scale bar is 0.5 mm.
(B) Representative adult plant photos of *ddm1 rdr2*, *ddm1 rdr2 rdr6* and *ccdr*.
(C) A silique photo of *ddm1 rdr2*, *ddm1 rdr2 rdr6* and *ccdr*.

### Figure S7

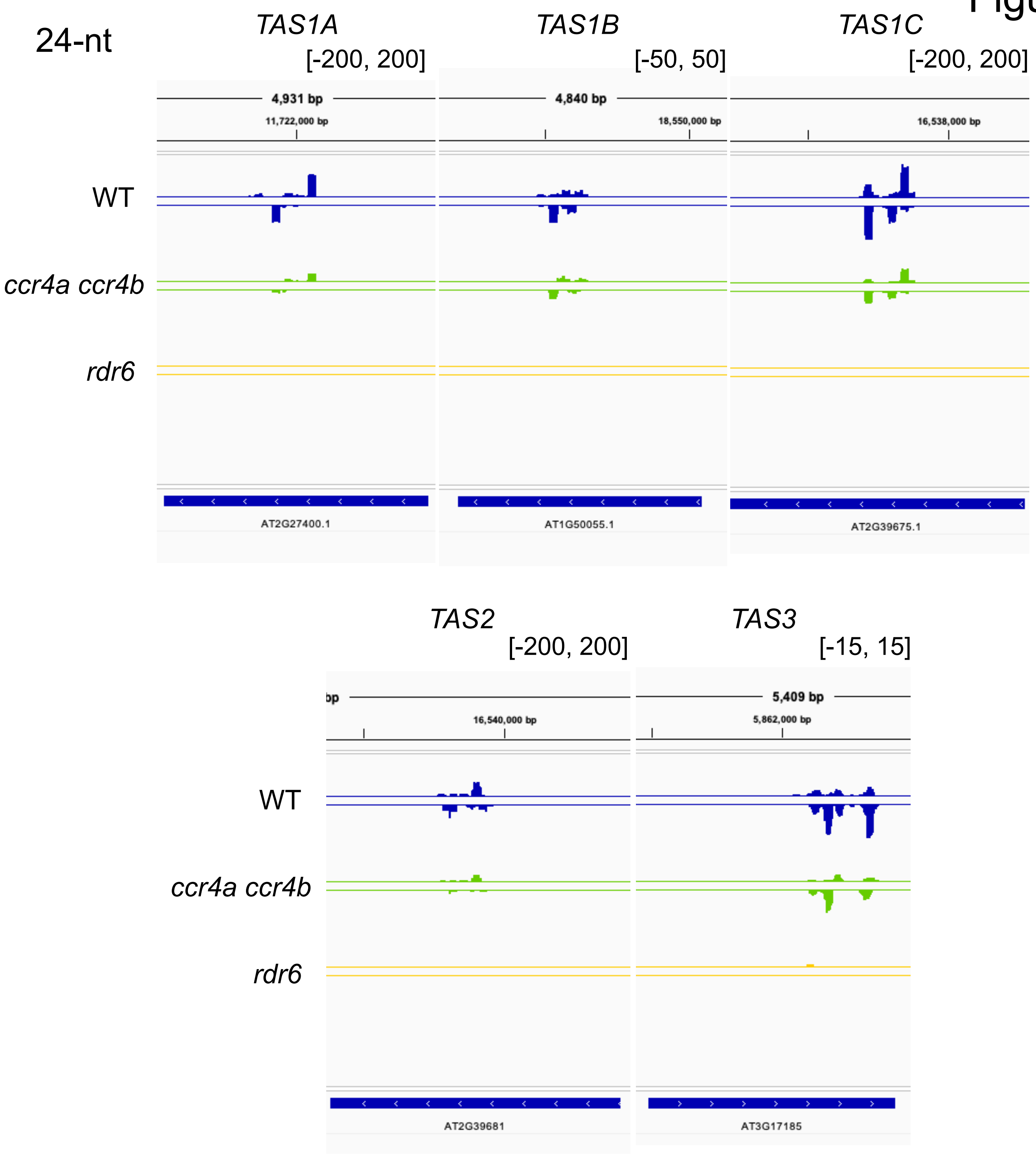


**Figure S7. 24-nt tasi-RNAs are reduced in *ccr4a ccr4b*, related to Figure 3.** Normalized 24-nt tasi-RNA levels (RPM) in WT, *ccr4a ccr4b* and *rdr6*.

Figure S8

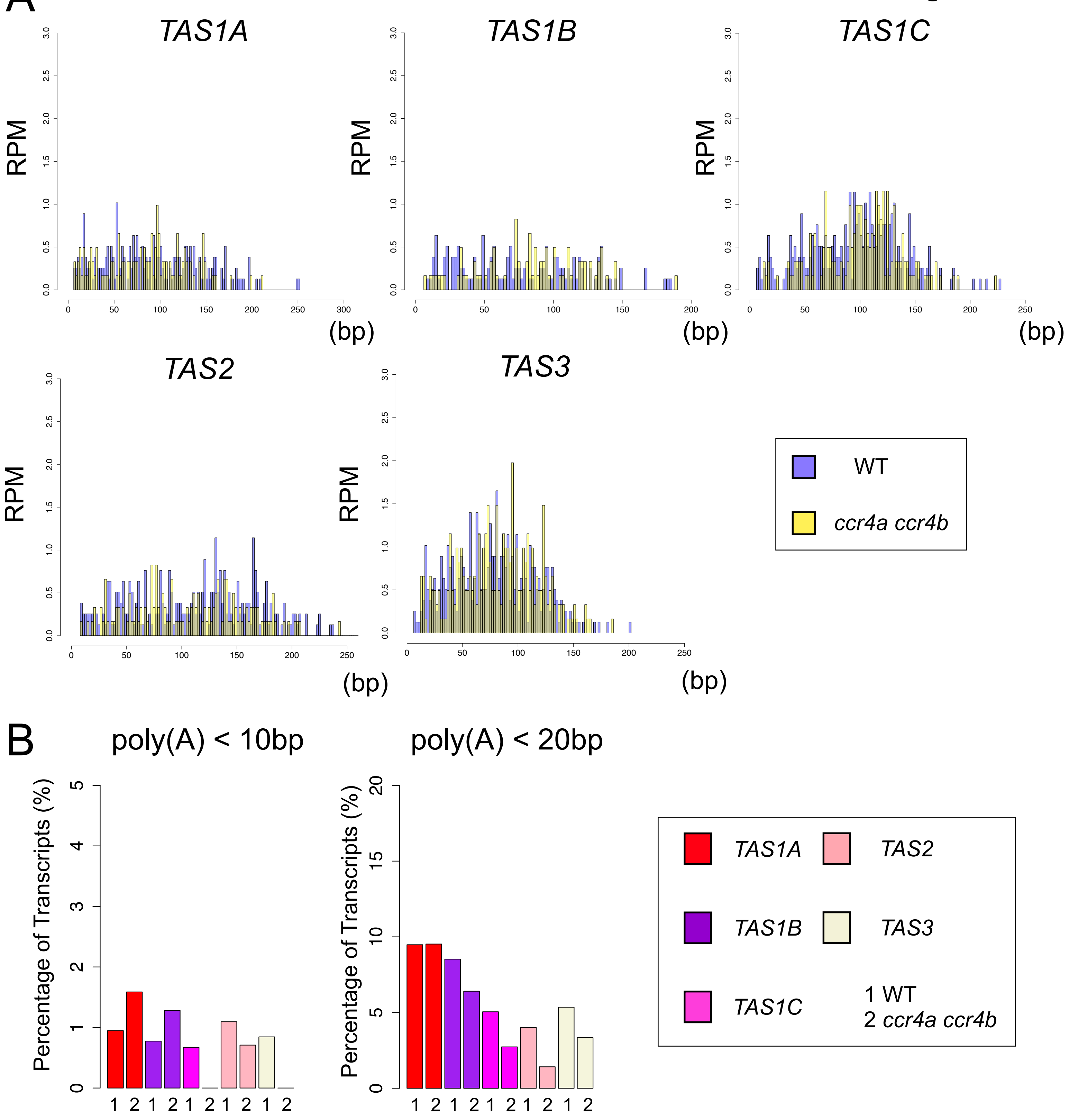


**Figure S8. Changes in poly(A) tail length of *TAS* RNAs in *ccr4a ccr4b*, related to Figure 4.**
(A) Distribution patterns of poly(A) tail length of *TAS* RNAs in WT (purple) and *ccr4a ccr4b* (yellow). Each bar in the histogram represents 2 bp bin.
(B) Proportion of short poly(A)-tailed transcripts from *TAS* loci in WT (1) and *ccr4a ccr4b* (2). Y-axis indicates the percentage (%) of ONT-DRS reads with short poly(A) tail (poly(A) < 10 bp or < 20 bp) relative to all reads mapped to the indicated *TAS* loci.

### Figure S9

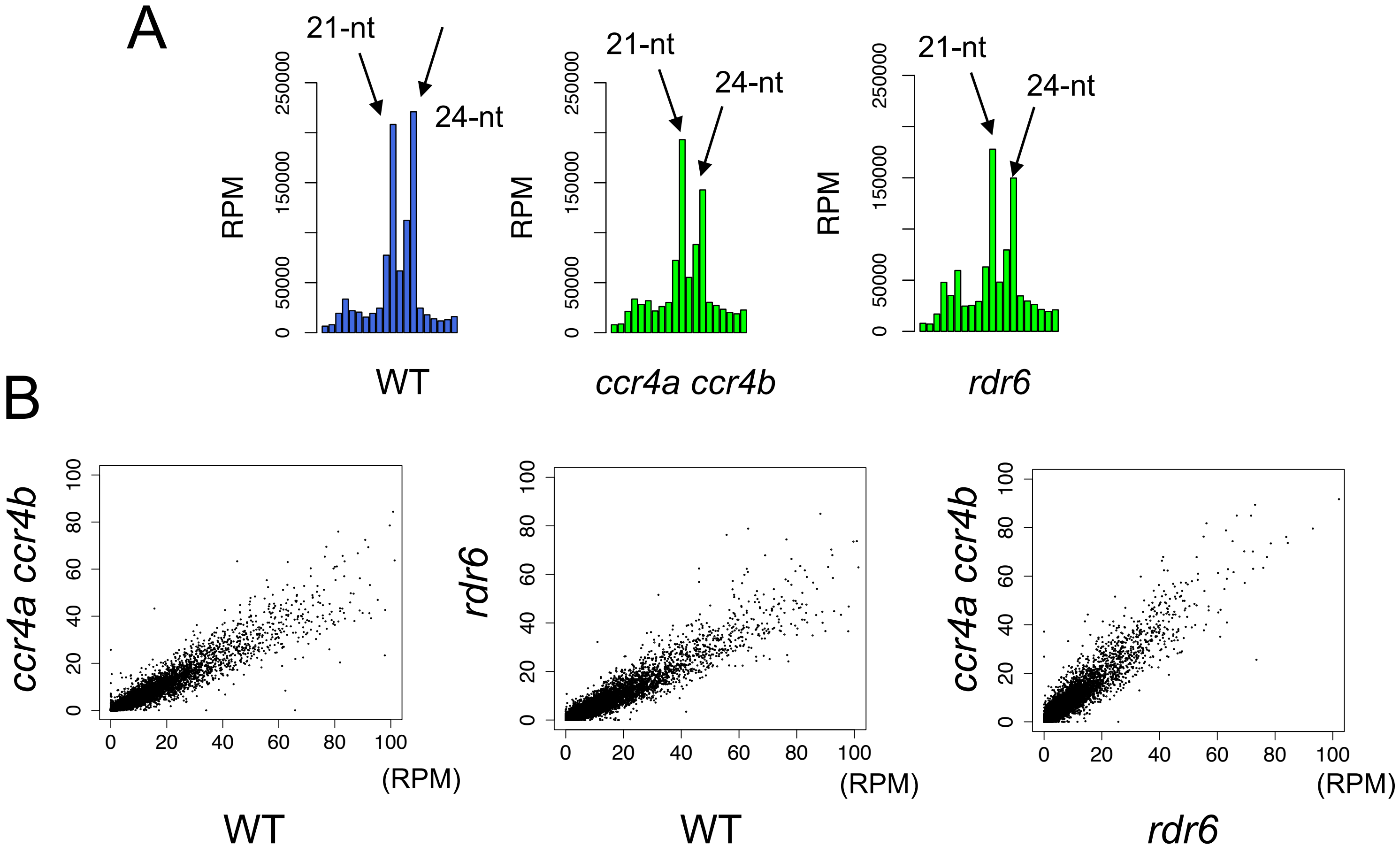


**Figure S9. 24-nt siRNAs are decreased in *ccr4a ccr4b* and *rdr6*, related to Figure 5.**
(A) Size distribution of small RNAs in WT, *ccr4a ccr4b* and *rdr6*. Barplots show normalized read count (RPM) from 11-nt to 30-nt small RNAs.
(B) Scatterplots comparing normalized 24-nt siRNA levels (RPM) at TEs across WT, *ccr4a ccr4b* and *rdr6*.

Figure S10

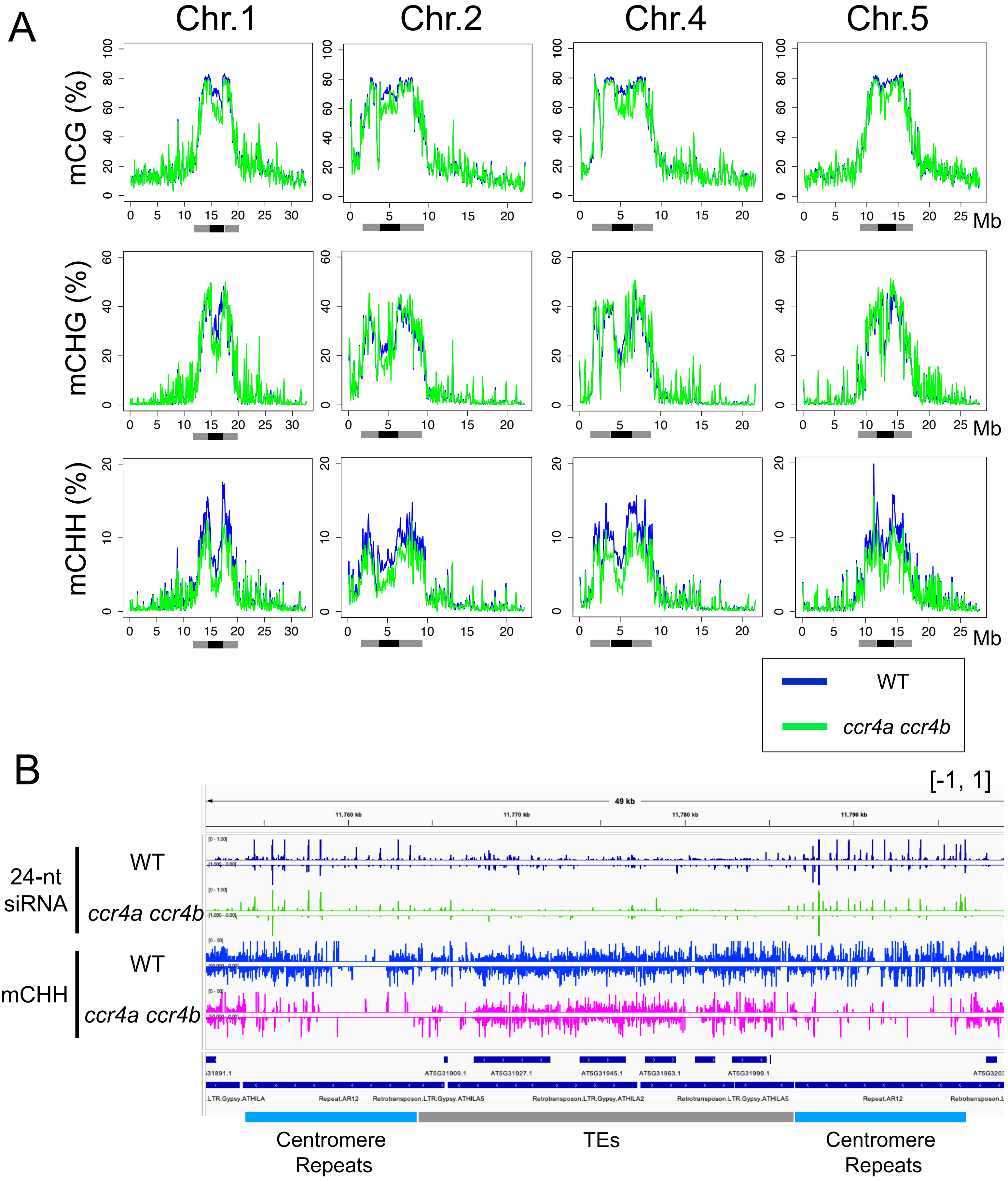


**Figure S10. Centromeric DNA methylation is reduced in *ccr4a ccr4b*, related to Figure 5.**
(A) Genome-wide DNA methylation levels at CG, CHG and CHH sites on chromosomes 1, 2, 4, 5 in WT (blue) and *ccr4a ccr4b* (green). Gray and black bars indicate pericentromeric regions and centromere repeat domains, respectively.
(B) 24-nt small RNA and DNA methylation (mCHH) tracks at the representative centromere region in WT and *ccr4a ccr4b*.

Figure S11

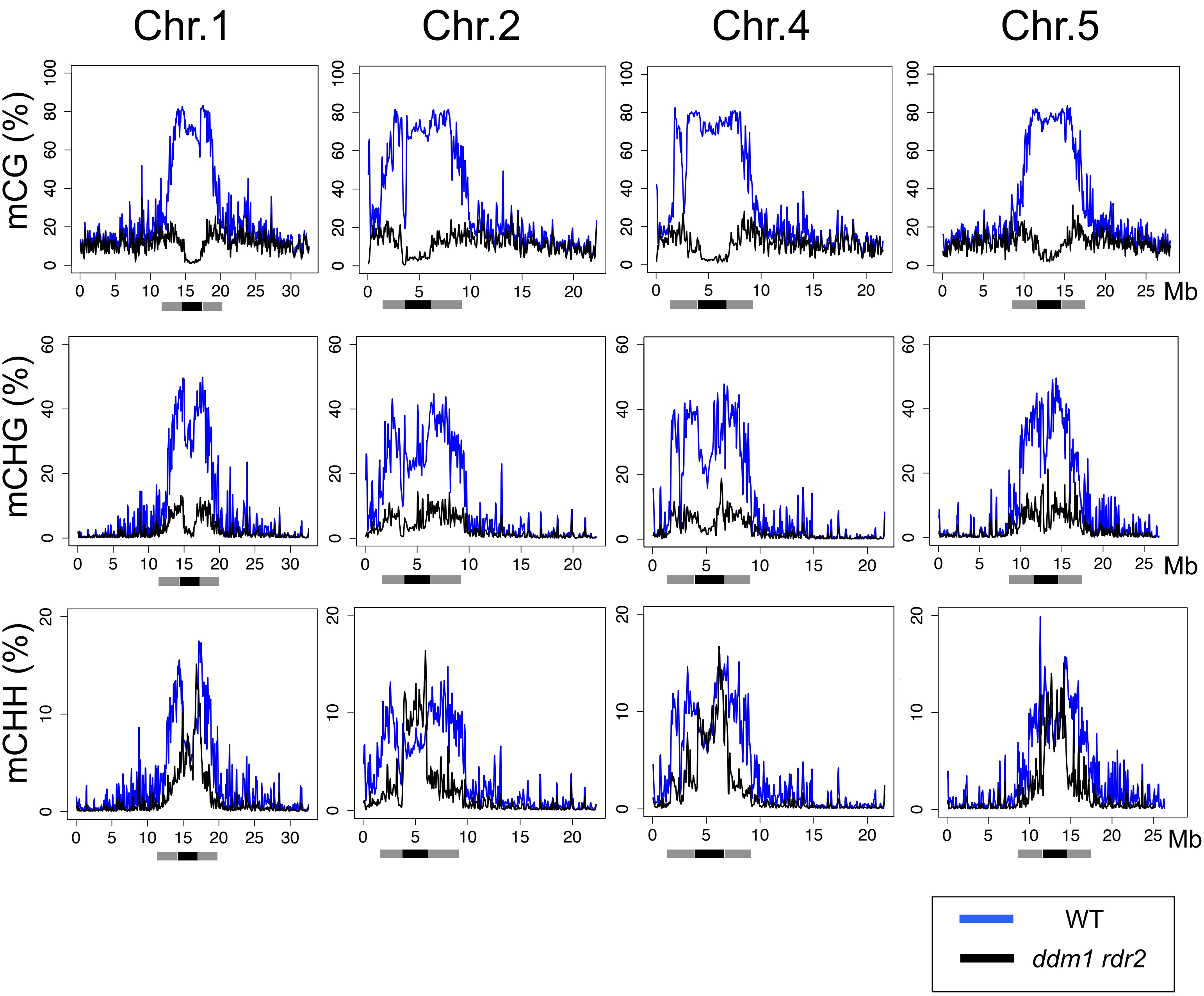


**Figure S11. CHH methylation at the centromere core is unaffected in *ddm1 rdr2*, related to Figure 5.**
Genome-wide DNA methylation levels at CG, CHG and CHH sites (Chr.1, 2, 4, 5) in WT (blue) and *ddm1 rdr2* (black). Gray and black bars indicate pericentromeric regions and centromere repeat domains, respectively.

Figure S12

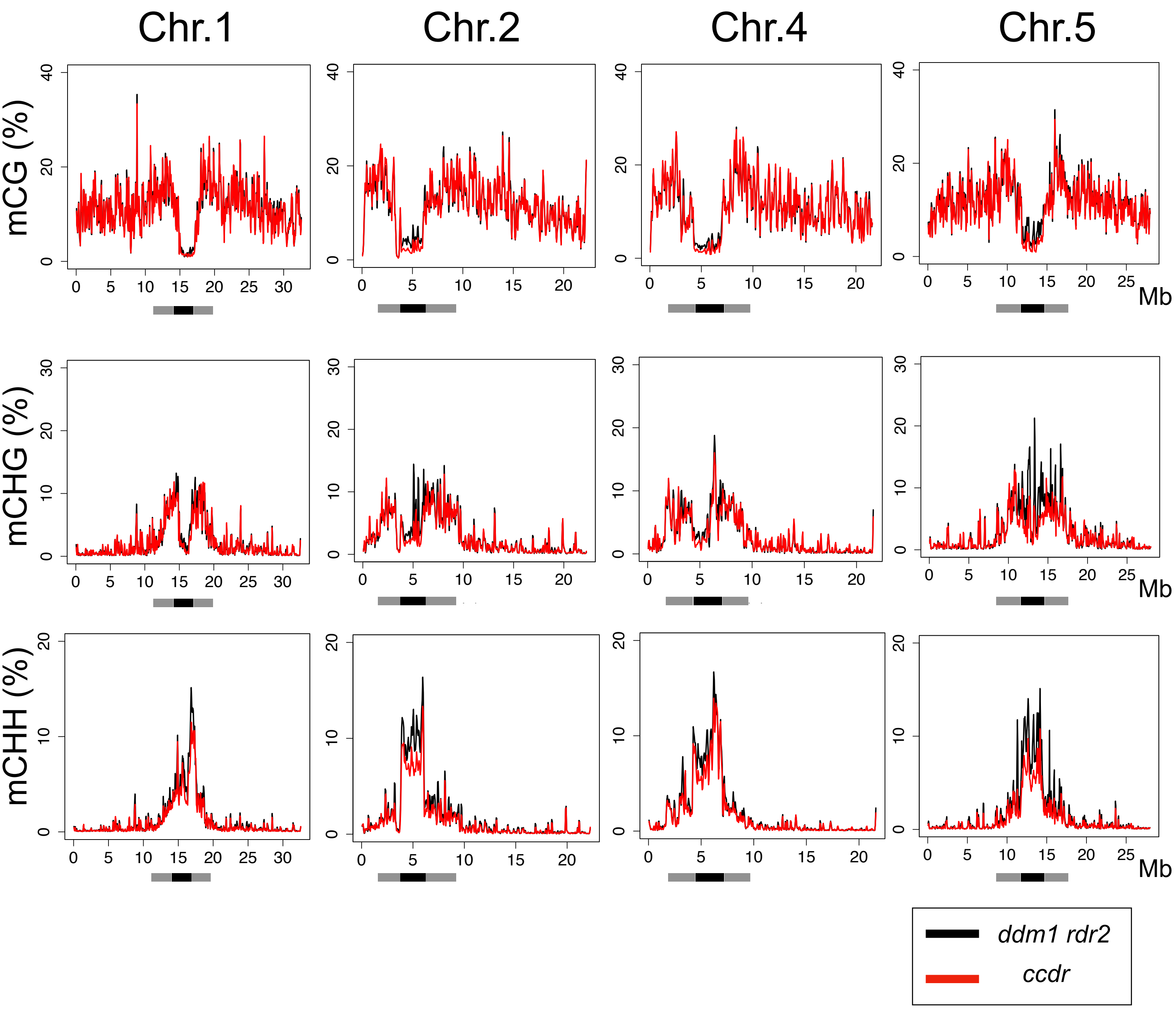


**Figure S12. *ccdr* exhibits further reduction of CHH methylation at the centromere core regions compared to *ddm1 rdr2*, related to Figure 5.**
Genome-wide DNA methylation levels at CG, CHG and CHH sites (Chr.1, 2, 4, 5) in *ddm1 rdr2* (black) and *ccdr* (red). Gray and black bars indicate pericentromeric regions and centromere repeat domains, respectively.

### Figure S13

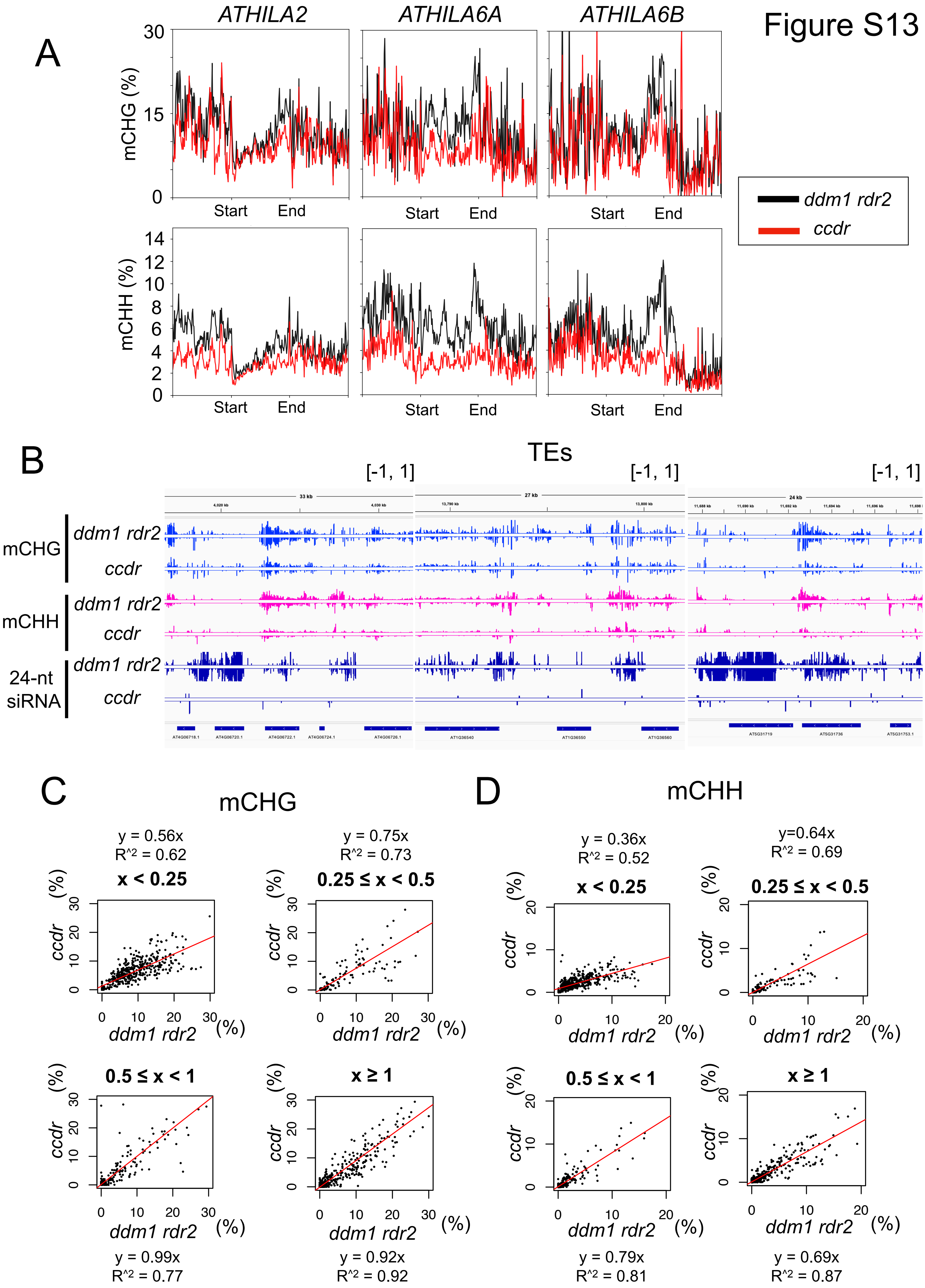
A
ATHILA2
ATHILA6A
ATHILA6B
mCHG (%)
mCHH (%)
Start
End
ddm1 rdr2
ccdr
B
TEs
[-1, 1]
mCHG
mCHH
24-nt siRNA
C
mCHG
D
mCHH
y = 0.56x
R^2 = 0.62
x < 0.25
y = 0.75x
R^2 = 0.73
0.25 ≤ x < 0.5
y = 0.36x
R^2 = 0.52
y=0.64x
R^2 = 0.69
0.5 ≤ x < 1
x ≥ 1
y = 0.99x
R^2 = 0.77
y = 0.92x
R^2 = 0.92
y = 0.79x
R^2 = 0.81
y = 0.69x
R^2 = 0.87

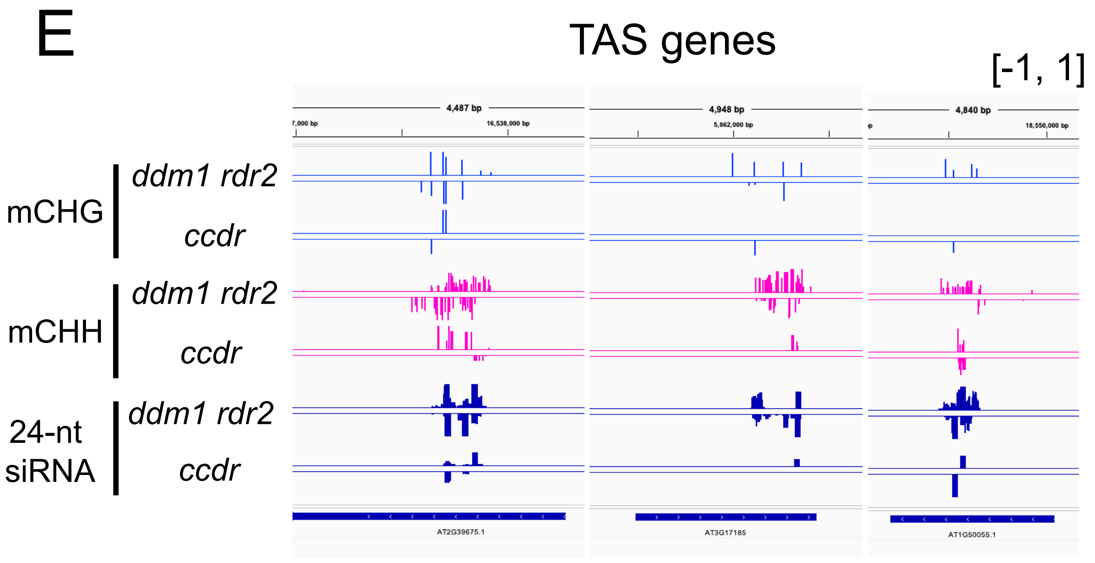


**Figure S13. CCR4-dependent siRNAs promote non-CG methylation at their target loci, related to Figure 5.**
(A) Metaplots of DNA methylation (mCHG and mCHH) over *ATHILA2*, *ATHILA6A* and *ATHILA6B* (> 4kb) in *ddm1 rdr2* (black) and *ccdr* (red).
(B) Non-CG DNA methylation (mCHG, mCHH) and 24-nt small RNA tracks over the regions associated with CCR4-dependent siRNAs.
(C-D) The effect of CCR4-dependent siRNAs on DNA methylation. The *Arabidopsis* genome sequence was binned into 4 kb and the bins were categorized depending on its associated CCR4-dependent siRNA levels. The bins associated with more than 0.15 RPM of 24-nt small RNAs were extracted, and these bins were divided into four categories based on the ratio (x) of normalized 24-nt small RNA levels in *ccdr* to those in *ddm1 rdr2*. Category 1; $x < 0.25$, Category 2; $0.5 > x \geq 0.25$, Category 3; $1 > x \geq 0.5$, Category 4; $x \geq 1$. mCHG and mCHH levels from each bin were plotted comparing between *ddm1 rdr2* and *ccdr* in each category. Regression line is represented with red line, and its equation and R-squared value is shown above or below the plot.
(E) Non-CG DNA methylation and 24-nt small RNA tracks at *TAS* genes in *ddm1 rdr2* and *ccdr*.

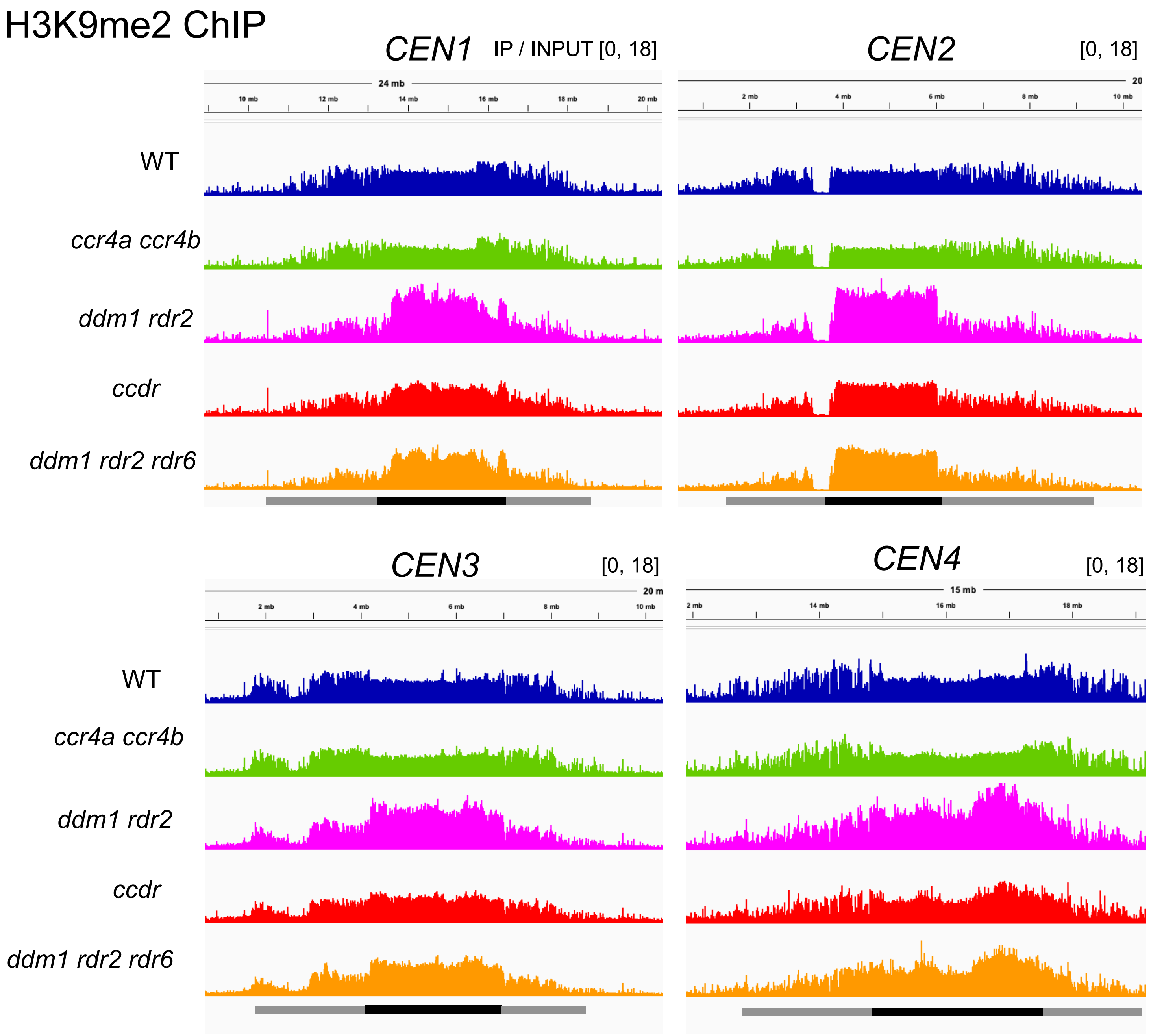


**Figure S14. The detailed centromeric H3K9 methylation landscape, related to Figure 6.** H3K9 methylation tracks over the centromere regions (*CEN1-4*). Gray and black bars indicate pericentromeric regions and centromere repeat domains, respectively.

### Figure S15

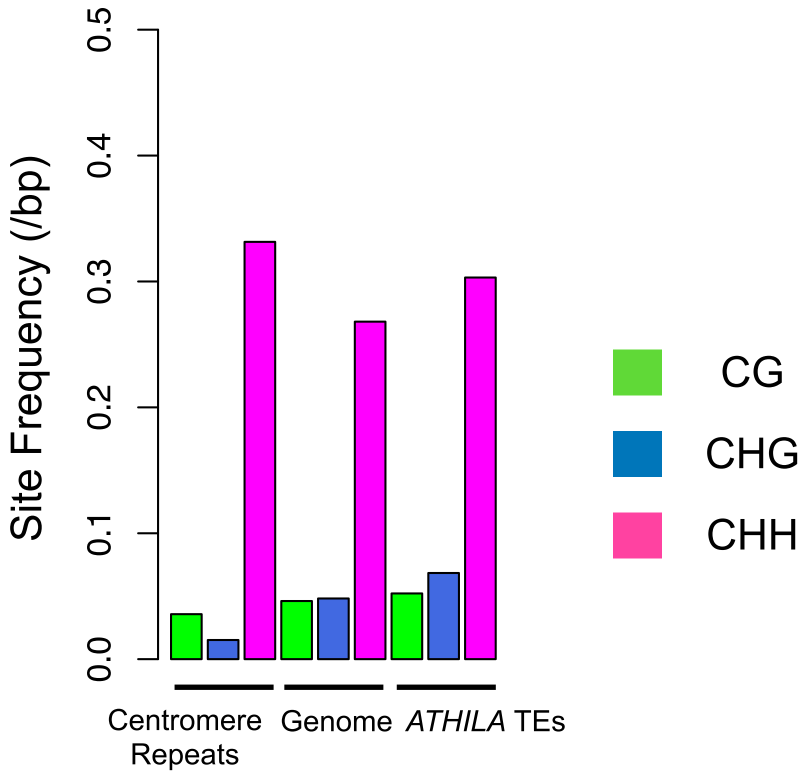


**Figure S15. The frequency of CHH site is higher in centromere repeats than those in the genome or *ATHILA* TEs, related to Figure 6.**
Comparison of cytosine context frequency (CG, CHG, CHH) observed in centromere repeats, the *Arabidopsis* whole genome and *ATHILA* TEs. Cytosine context frequency was calculated by normalizing the number of cytosines in each context with the chromosome size (bp) of each category.

Figure S16

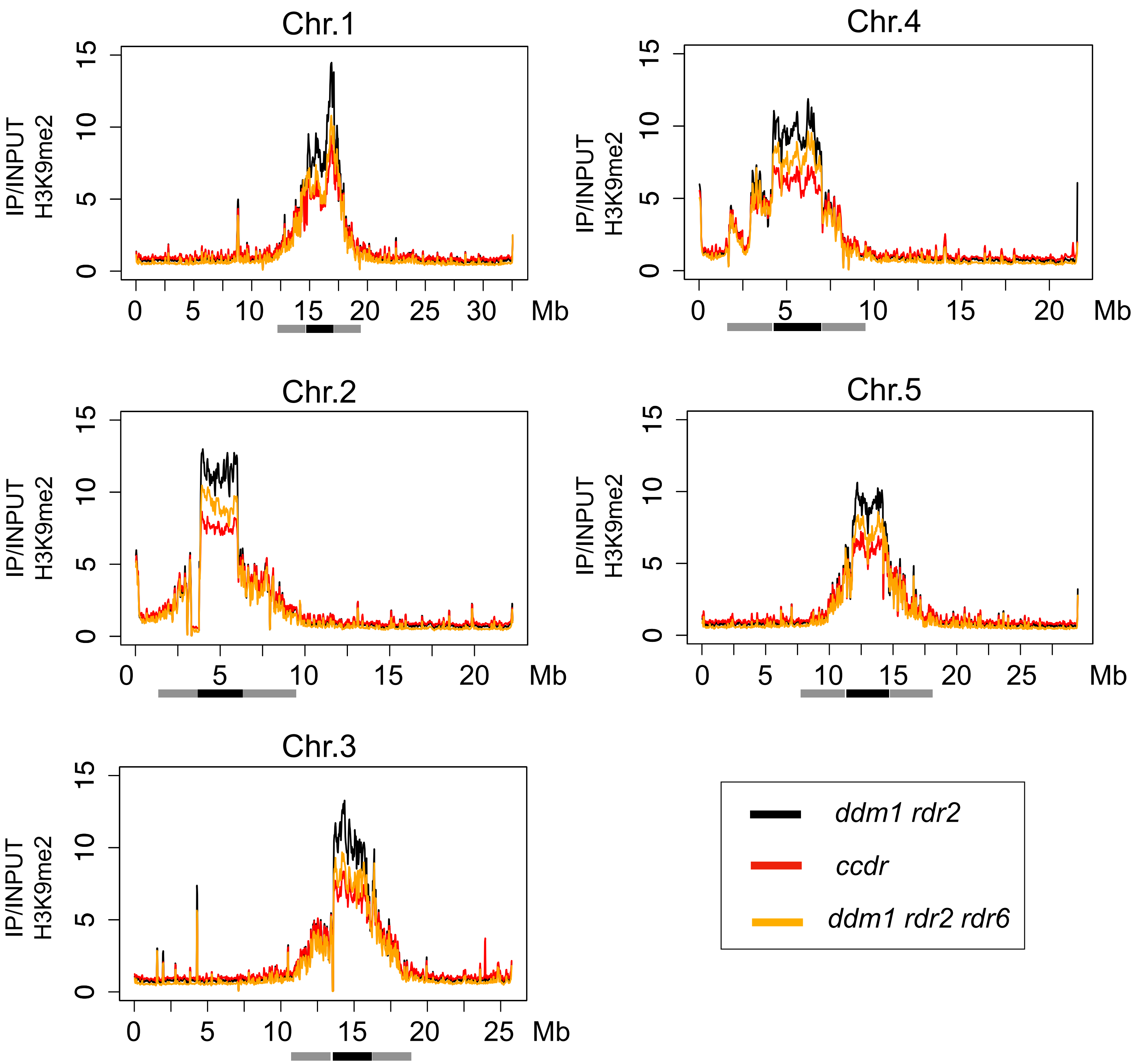


**Figure S16. CCR4 and RDR6 maintain centromeric H3K9 methylation in *ddm1 rdr2*, related to Figure 6.**
Genome-wide H3K9 methylation signals (IP/INPUT) in a 50 kb sliding window comparing across *ddm1 rdr2* (black), *ddm1 rdr2 rdr6* (orange) and *ccdr* (red). Gray and black bars indicate pericentromeric regions and centromere repeat domains, respectively.

| Gene_ID | WT (RPM) | *ccr4a ccr4b* (RPM) | *ccr4a ccr4b* / WT | Gene_name |
|---|---|---|---|---|
| AT3G17185 | 3330.2 | 942.5 | 0.28 | *TAS3* |
| AT2G27400 | 4016.4 | 1834.1 | 0.46 | *TAS1A* |
| AT2G39681 | 10188.5 | 5010.7 | 0.49 | *TAS2* |
| AT2G39675 | 9239.4 | 5280.9 | 0.57 | *TAS1C* |
| AT1G50055 | 1724.7 | 1025.8 | 0.59 | *TAS1B* |
| AT5G49615 | 24.3 | 5.1 | 0.21 | *TAS3B* |

**Table S1. Mutation of *CCR4* down-regulates tasiRNA production, related to Figure 2.**

Normalized 21-nt small RNA levels (RPM) of *TAS* genes in *ccr4a ccr4b* comparing to WT.

| Gene Name | Fold Change (Log2 *ccdr* /*ddm1 rdr2*) |
|---|---|
| *SGS3* | 0.164 |
| *AGO7* | -0.235 |
| *SDE5* | 0.165 |
| *DCL4* | -0.073 |
| *RDR6* | -0.082 |

**Table S2. *CCR4* mutation does not impact the expression of genes involved in RDR6-dependent siRNA synthesis, related to Figure 3.**Normalized RNA-seq read count (CPM) from genes involved in the RDR6-dependent siRNA synthesis pathway were compared between *ddm1 rdr2* and *ccdr*. The fold changes were calculated based on the data from three biological replicates of RNA-seq data.

| | | Reads containing centromere repeat sequence | Total mapped reads |
|---|---|---|---|
| WT | Rep 1 | 246 | 16489606 |
| | Rep 2 | 290 | 15058985 |
| | Rep 3 | 417 | 17842316 |

**Table S3. RNA-seq reads containing centromere repeat sequence are present in WT, related to Figure 5.**

The number of mapped reads containing centromere repeat sequence in WT RNA-seq data (three biological replicates).

**mCHG**

| Down | Total | % |
|---|---|---|
| 320 | 479 | 66.8 |
| 100 | 170 | 58.8 |
| 188 | 390 | 48.2 |
| 377 | 736 | 51.2 |

**mCHH**

| Down | Total | % |
|---|---|---|
| 326 | 479 | 68.0 |
| 108 | 168 | 63.5 |
| 198 | 374 | 50.7 |
| 434 | 652 | 58.9 |

**Table S4. Loss of CCR4-dependent siRNAs causes reduction of non-CG methylation at their target loci, related to Figure S13.**The number of total bins from each category used in Figures S13C and S13D are shown (Total). Of these bins, the number of bins that showed reduced mCHG and mCHH in *ccdr* compared to *ddm1 rdr2* was counted (Down), and its percentage (%) was calculated.

| | |
|---|---|
| CCR4B HP up apa1 fw | AAAGGGCCCACAGAGTGATAAGAAAGTAA |
| CCR4B HP up rv | CCGACGCGAAGCGGGTAGATTTCTTGCAGGCAAACTACAT |
| CCR4B HP down spe1 fw | AAAACTAGTACAGAGTGATAAGAAAGTAA |
| CCR4B HP down xho1 rv | AAACTCGAGTTCTTGCAGGCAAACTACAT |
| GUS spacer fw | ATCTACCCGCTTCGCGTCGG |
| GUS spacer xho1 rv | ATTCTCGAGCGAGTGAAGATCCCTTTCTT |

**Table S5. Oligonucleotides used to generate the plasmid for *CCR4B* knock down, related to Figure 1.**
